# Marked sex differences and frequent carriage of recognized uropathogens in the human urogenital microbiome: urine profiles from a population-level shotgun metagenomics study

**DOI:** 10.64898/2026.02.25.707997

**Authors:** Tiscar Graells, Yi-Ting Lin, Paula Rodríguez García, Oksana Lukjancenko, Tessa Schillemans, Janne Marie Moll, Henrik Bjørn Nielsen, Shafqat Ahmad, Tove Fall, Johan Ärnlöv

## Abstract

**Background:** The urinary tract was long considered sterile. The discovery of the urinary microbiome has attracted interest in its role in urinary health and disease. Population-level urinary microbiome composition remains poorly defined and previous studies relied on selected populations and low-resolution sequencing. We characterized the microbial composition of voided urine from adults without urinary symptoms in a large Swedish population cohort to establish a population-based reference for microorganisms detected in voided urine using shotgun metagenomics.

**Methods:** We performed shotgun metagenomic sequencing of voided urine samples from 2,062 participants of the population-based Swedish CArdioPulmonary bioImage Study (SCAPIS; 55% women, 50-65 years).

**Results:** Among 3,668 identified species, 90% were classified at the species level. Microbial profiles differed substantially between sexes, with lactobacilli and bifidobacteria enriched in females, while *Cutibacterium acnes, Enterococcus* and *Propionimicrobium lymphophilum* were enriched in males. Recognized uropathogens, including *Enterococcus faecalis* and *Escherichia coli*, were frequently detected despite participants having no urinary symptoms. Males showed higher alpha diversity (Shannon and inverse Simpson indices), whereas richness did not differ by sex. Age showed only modest associations with alpha diversity, and beta diversity differed by sex but showed minimal age-related variation.

**Conclusions:** This study provides a population-based reference for interpreting microorganisms detected in routine voided urine samples. Biological sex was the principal determinant of microbial composition, supporting routine consideration of sex in future studies of the urogenital microbiota using routine voided urine samples.

## 1. Introduction

Accurate interpretation of microorganisms detected in urine is fundamental to the diagnosis and management of urinary tract disease. Advances in culture-independent sequencing have demonstrated that urine contains microbial communities, changing the longstanding assumption that urine and the urinary tract were sterile [1,2]. Despite advances in human microbiome research, particularly of the gut microbiome, the urinary microbiome remains the least characterized human microbiome, partly due to its low microbial biomass and because it was discovered a bit more than a decade ago [1,2].

Most studies have investigated urine microbiota in patients with urinary tract infections (UTIs) or other urological disorders rather than healthy populations. Historically, urine microbiology has focused on UTIs using culture-based approaches, more recently including also expanded quantitative urine cultures [2,3]. UTIs are often associated with a dysbiotic microbiome composition, although the eubiotic (‘healthy’) urinary microbiome remains insufficiently characterized [1]. Consequently, the composition of the baseline symptom-free urinary microbiome remains poorly defined, limiting interpretation of microorganisms detected in urine from individuals without urinary symptoms.

As molecular methods become more used to characterize urinary microorganisms, defining the microbial composition of urine from individuals without urinary symptoms has become increasingly important for interpreting findings in both clinical and research settings [4]. Previous studies suggest marked sex differences in urinary microbial composition [1,5,6]; with lactobacilli frequently associated with urinary health and influenced by oestrogen in females [1,5,7,8]. The female urinary microbiome partially overlaps with the vaginal microbiome but these two represent distinct microbial niches which are not interchangeable [9,10]. Knowledge of the male urinary microbiome remains even more limited, with most studies focusing on patients with urological disease or studying the distal genitourinary tract [1,11,12].

Previous studies relied on culturomics or metataxonomics (16S/18S rRNA gene sequencing) and included relatively small, selected cohorts, predominantly females [13–15]. Although suprapubic aspiration and catheterization provide bladder-specific samples, these invasive collection methods are not common. In clinical practice and in most studies, the standard collection method is the voided midstream urine which is non-invasive and can be self-collected [4]. From a microbiological perspective, these urine samples may contain microbiota from the periurethral skin or genital tract [16]. However, the contribution of microorganisms originating from adjacent anatomical sites to voided urine profiles remains also incompletely understood.

Without a robust population reference describing microbes in urine from asymptomatic adults, the clinical significance of microorganisms detected remains uncertain. This limits the ability to distinguish microbial carriage from findings that may reflect urinary tract disease. This study was designed to characterize microbial profiles in voided urine representative of routine clinical urine collection from adults free of urinary symptoms. The objective was also to establish a population reference of the microbial composition of voided urine while characterizing sex and age associated differences. Therefore, we performed high resolution shotgun metagenomic sequencing of voided urine of more than 2,000 participants from the population-based Swedish Cardiopulmonary bioimage study (SCAPIS) cohort [17].

## 2. Materials and methods

### 2.1 Cohort description and ethics

SCAPIS is a Swedish multicentre population-based cohort with approximately 5,000 individuals aged 50-65 years per location (Gothenburg, Linköping, Malmö/Lund, Stockholm, Umeå and Uppsala) designed to improve understanding of cardiovascular and pulmonary disease while providing a well-characterized population for epidemiological research [17]. Participants were recruited from the general population and were not selected based on diseases, nor urinary symptoms or urological disease. SCAPIS was approved by the ethics committee at Umeå University (DNR 2010-228-31M) and conducted in accordance with the Declaration of Helsinki. All participants provided written informed consent, and the present study was approved by the Swedish Ethical Review Authority (DNR 2018-315; SCAPIS petition 210).

### 2.2 Participants and sample analyses

This SCAPIS study includes participants from the SCAPIS-Uppsala (n=2,062), data at inclusion included samples from visits to a test centre or collected as part of a core questionnaire (e.g. age, sex, smoking status, alcohol intake, diet). We conducted a cross-sectional analysis of voided urine samples collected from participants in the SCAPIS-Uppsala cohort. Participants provided their midstream voided urine in the morning when they came to the clinic, and previously they were advised to use the clean-catch method. Participants were instructed to attend the study visit only if they were free of acute illness or infection at the time of sample collection. Samples were aliquoted in tubes (11 mL) and immediately frozen. Because voided urine may contain microorganisms originating from the lower urinary tract, periurethral skin and genital tract, these samples were considered representative of the urogenital microbiota [16].

Samples were shipped frozen and in dry ice to ensure preservation in two batches to Cmbio (Copenhagen, Denmark). The first batch comprised 1,000 samples, while the second batch consisted of 1,062 urine samples and, because SCAPIS is a population-based study, the age range of the cohort (50–65 years) includes individuals with conditions such as diabetes, kidney disease and previous cardiovascular events among other comorbidities **(Table 1** and **Table S1)**. DNA was extracted and shotgun sequenced at Cmbio. Briefly, urine was centrifuged, and pellets were used for DNA extraction using DNeasy 96 Blood & Tissue kit (Qiagen, Handbook version 07/2020) with lysozyme and proteinase K treatments. Three negative controls and one positive control (ZymoBIOMICS™ Microbial Community Standard, Zymo Research) were included per batch of samples during DNA extraction. Genomic DNA was fragmented (≈ 350 bp) and libraries were constructed using NEBNext Ultra Library Prep for Illumina (New England Biolabs). DNA libraries were sequenced using 2 × 150 bp paired-end sequencing on a NovaSeq 6000 Illumina platform (Illumina, USA). On average, 37.2 million read pairs and 32.7 million read pairs per sample were obtained in the first batch and the second batch, respectively, with a combined average of 34.9 million read pairs. Shotgun metagenomic reads were quality filtered and host-derived sequences were removed by alignment against the human reference genome GRCh38. On average 67% of reads per sample were identified as host-derived and removed. The remaining reads were defined as high-quality non-host (HQNH) reads, with an average of 11.4 million read pairs per sample. HQNH reads were taxonomically profiled using the CHAMP™ pipeline with the HMR05 reference database [18]. Taxonomic annotation was assigned according to the Genome Taxonomy Database (GTDB) using a ≥95% average nucleotide identity threshold for species. On average, 7.8 million read pairs per sample mapped to the gene catalogue, representing 57.3% of HQNH reads.

**Table 1.** Descriptive table of our cohort and by biological sex. N = Number of samples. SD = standard deviation

|  | ALL (N = 1615) | FEMALES (N=891) | MALES (N=724) |
| --- | --- | --- | --- |
| **Age in years (mean (SD))** | 57.9 (4.4) | 57.6 (4.4) | 58.2 (4.4) |
| **Batch 1** | 800 (49.5%) | 438 (49.2%) | 362 (50.0%) |
| **BMI category** |  |  |  |
| Underweight | 6 (0.4%) | 6 (0.7%) | 0 (0%) |
| Standard weight | 509 (31.5%) | 341 (38.3%) | 168 (23.2%) |
| Overweight | 716 (44.3%) | 345 (38.7%) | 371 (51.2%) |
| Obesity | 373 (23.1%) | 195 (21.9%) | 178 (24.6%) |
| **Smoker** |  |  |  |
| Current smoker | 138 (8.9%) | 67 (7.7%) | 71 (10.3%) |
| Ex-smoker | 518 (33.2%) | 324 (37.3%) | 194 (28.1%) |
| Never | 880 (54.4%) | 468 (53.9%) | 412 (59.7%) |
| **Alcohol consumption** |  |  |  |
| Never | 133 (8.24%) | 83 (9.32%) | 50 (6.91%) |
| Occasionally, less than once a month | 255 (15.8%) | 160 (18.0%) | 95 (13.1%) |
| 2-4 times/month | 580 (35.9%) | 326 (36.6%) | 254 (35.1%) |
| 2-3 times/week | 476 (29.5%) | 257 (28.8%) | 219 (30.2%) |
| Regularly, 4 or more times a week | 99 (6.13%) | 37 (4.15%) | 62 (8.56%) |
| **Physical Activity** |  |  |  |
| Mainly sedentary | 328 (20.3%) | 200 (22.4%) | 128 (17.7%) |
| Light intensity | 296 (18.3%) | 162 (18.2%) | 134 (18.5%) |
| Regular moderate intensity | 574 (35.5%) | 320 (35.9%) | 254 (35.1%) |
| Regular high intensity | 338 (20.9%) | 172 (19.3%) | 166 (22.9%) |
| **Education** |  |  |  |
| Incomplete compulsory | 13 (0.80%) | 7 (0.79%) | 6 (0.83%) |
| Compulsory | 109 (6.75%) | 44 (4.94%) | 65 (8.98%) |
| Upper secondary | 660 (40.9%) | 344 (38.6%) | 316 (43.6%) |
| University | 771 (47.7%) | 469 (52.6%) | 302 (41.7%) |
| **Birthplace** |  |  |  |
| Sweden | 1332 (82.5%) | 744 (85.6%) | 588 (85.2%) |
| **Diabetes** | 204 (12.6%) | 93 (10.4%) | 111 (15.3%) |
| **Chronic kidney disease** |  |  |  |
| None or stage 1 | 760 (47.1%) | 412 (46.2%) | 348 (48.1%) |
| Stage 2 | 803 (49.7%) | 451 (50.6%) | 352 (48.6%) |
| Stage 3 | 47 (2.9%) | 26 (2.9%) | 21 (2.9%) |
| Stage 4 | 2 (0.1%) | 1 (0.1%) | 1 (0.1%) |
| **Previous cardiovascular events (questionnaire)** | 141 (8.7%) | 50 (5.6%) | 91 (12.6%) |
| **Subclinical atherosclerosis**<br>**(by Coronary Artery Calcium score)** |  |  |  |
| None | 947 (58.6%) | 632 (70.9%) | 315 (43.5%) |
| Mild | 403 (25%) | 186 (20.9%) | 217 (30%) |
| Moderate | 151 (9.3%) | 47 (5.3%) | 104 (14.4%) |
| Severe | 114 (7%) | 26 (2.9%) | 88 (12.2%) |

Background species signals that likely originate from amplification of reagent contaminants, environmental contaminants, or other background sources were identified as follows. First, species were flagged as potential background signal if (a) the species was detected in at least ten samples and there was a Pearson correlation coefficient < −0.80 between the species relative abundance and the sample’s sequencing depth; or (b) groups of species present in at least ten samples had a co-abundance correlation coefficient > 0.90 among each other [19]. Species identified as potential background signal were removed from further analysis if they fulfilled any of the following additional criteria: (a) the species is a typical reagent contaminant [19]; (b) typical environmental or otherwise non-human associated species; or (c) unexpectedly observed in positive controls. In total, 201 species were identified as background signal and removed from the dataset before any additional analysis.

### 2.3 Exclusion criteria

We excluded urine samples which failed DNA extraction, DNA sequencing (n=121) or lacked sequence reads mapped to the reference catalogue (n=2, n=121 and n=2, respectively) **(Figure 1)**. We further excluded one participant with two samples (duplicates). Samples with fewer than 10,000 microbial counts were also excluded from primary analyses (n=320) to ensure robust estimation of microbial community composition in this low-biomass sample type **(Figure 1)**. Finally, 1,615 urine samples from unique participants were included in our primary analyses, comprising 724 from males and 891 from females.

**Figure 1:**
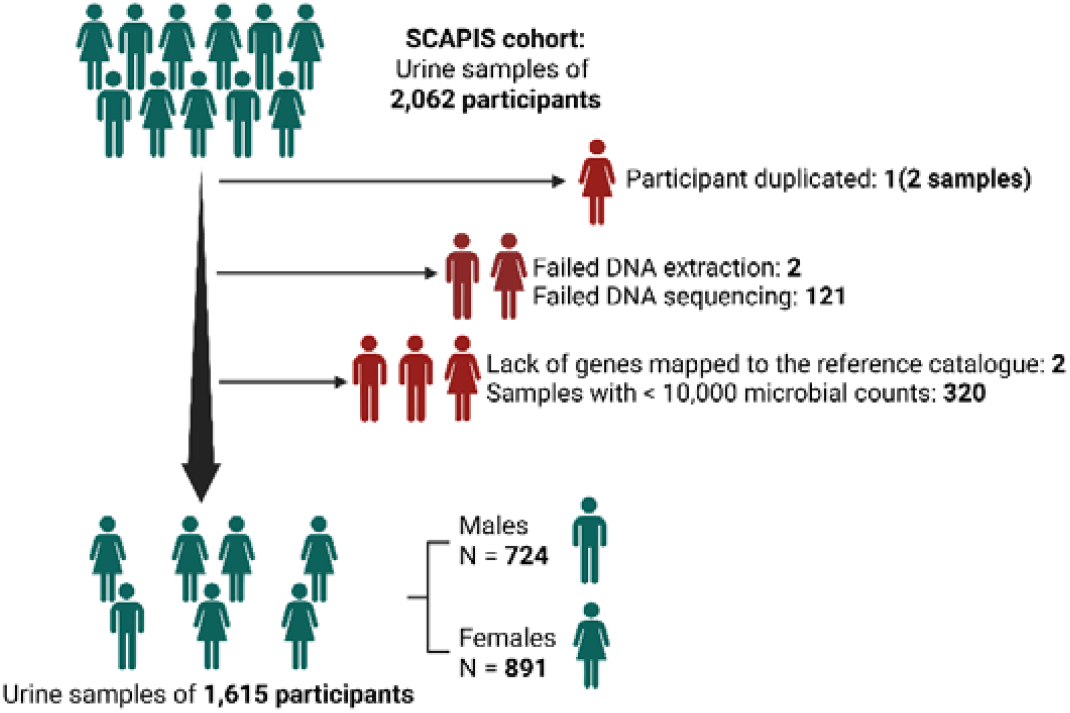
Flowchart of urine samples and participants in the study.

### 2.4. Statistical analyses

All statistical analyses were performed using R (version 4.4.2). Differences in sequencing depth were addressed by rarefying samples to 10,000 microbial reads prior to estimation of alpha diversity indices. Diversity indices were estimated with the R packages vegan (version 2.7-1). Primary analyses focused on differences according to sex and age because these variables were specified a priori as the principal biological factors of interest. Associations between alpha diversity (richness, Shannon, and inverse Simpson) and sex and age group (50-55, 56-60, 61-65 years) were assessed using regression models adjusting for batch. Richness was analysed using negative binomial models to account for overdispersion, while Shannon and inverse Simpson indices were analysed using linear models. All models included sex, age group, and their interaction term. P-values were adjusted using the Benjamini-Hochberg false discovery rate (FDR).

Beta diversity indices, including Bray-Curtis dissimilarity, Jaccard, and Sørensen indices were computed using relative abundance data derived from raw counts, which reduces the influence of differences in sequencing depth between samples. We tested differences in beta diversity for sex and age by Permutational Multivariate Analysis of Variance (permutation test for adonis under reduced model settings: 999 permutations, free). Beta diversity was plotted using principal coordinates analysis (PCoA) using the package ape (version 5.8-1). Homogeneity of group dispersions was evaluated using permutation-based tests with Tukey post hoc comparisons. To assess whether sex was driving these differences, we performed a constrained ordination PCoA to restrict the axes to variation explained by our predictor variable (sex) using the function ‘capscale’ in the vegan package.

For sensitivity analyses of alpha and beta diversity indices, samples with more than 1,000 microbial counts were also included to retain very low biomass samples (n = 256 (samples with >1000 and < 10,000 microbial counts); total of 1871 urine samples). Sensitivity analyses for alpha diversity indices were performed using a lower rarefaction threshold of 1,000 reads per sample, while for beta diversity indices these were conducted to evaluate robustness to sequencing depth.

Microbial composition was checked at different taxonomical levels to look for differences between sexes. For specific analyses, we applied prevalence and rarity filters to reduce noise and limit false-positive results. We retained species present in at least 5% of female or male samples, or in at least 5% of samples across the entire cohort. In addition, we kept species that had a minimum of 500 sequence reads in five or more samples to retain rare but occasionally abundant species.

We performed two exploratory analyses that do not allow for covariate adjustment: (1) an indicator species analysis to detect taxa that are strongly associated with each specific sex using the package indicspecies (version 1.8.0) with BH FDR correction; and (2) a linear discriminant analysis effect size (LefSe) with FDR correction with microbiomeMarker (version 1.12.2) with Kruskal-Wallis tests and linear discriminant analyses score higher than 2 for effect size estimation. LEfSe was run using cumulative sum scaling normalization that stabilizes variance across samples, which is robust for sparse microbiome data and taxa filtering [20]. We also performed three differential abundance analyses using counts with statistical batch correction. The first analysis was performed using DESeq2, while the second analysis accounted for sample variability and was evaluated using ALDEx2 (version 1.38.0); both analyses applied BH FDR correction. The third analysis comprised microbiome differential abundance and correlation analyses with bias correction using ANCOM-BC (version 2.8.1), for which a conservative Holm correction was applied to control the family-wise error rate. ANCOM-BC and ALDEx2 are compositional data analysis methods that explicitly account for the relative nature of microbiome sequencing data. Spearman correlations were used to assess the consistency of effect sizes across the three methods. For these analyses, we performed a sensitivity analysis including age as a covariate to ensure that our conclusions were independent of age, given the 15-year age range among study participants. To address this, we tested for differences in age between sexes using a t-test, assessed multicollinearity between sex and age with Pearson correlation, and calculated the variance inflation factor (VIF) using the R package car (version 3.1.3). We also checked age with these three differential abundance analyses to study the influence of age in the urogenital microbiome.

We applied three supervised machine learning approaches to predict sex from urinary microbiome profiles. For that, we used R packages caret (version 7.0-1), caretEnsemble (version 4.0.1) and pROC (version 1.18.5), while specific random forest, LASSO logistic regression and gradient boosting were evaluated by R packages randomForest (version 4.7-1.2), glmnet (version 4.1-10) and xgboost (version 1.7.11.1), respectively. We used normalized count data controlling for batch and age as covariates. We implemented 5-fold cross-validation stratified by batch–sex combinations, which ensured that samples from the same batch were not split between training and validation sets and, thus, avoid batch leakage between training and validation sets. Then, random forest classifiers were trained to assess feature importance via mean decrease in Gini, while LASSO logistic regression (α=1) was fitted to identify predictive taxa, and gradient boosting was trained to capture non-linear relationships. Model performances were evaluated by cross-validated area under the ROC curve (AUC), sensitivity, and specificity. In addition, we did a permutation test shuffling labels to test if AUC high values were dropping to confirm that our initial AUC results were driven by sex and not capturing artifacts.

## 3. Results

### 3.1 Descriptive characteristics

From the original urine samples, 1,615 were included in the final analysis **(Figure 1)**. Baseline characteristics of participants are described in **Table 1**. The mean age was 57.9 years, and 55.8% of participants were female. Males were more likely than females to be overweight or obese, to smoke, and to consume alcohol, had lower education attainment, exhibited a higher prevalence of diabetes, previous cardiovascular events and subclinical atherosclerosis **(Table 1)**.

### 3.2 Prevalence of microorganisms and recognized uropathogens in voided urine

Recognized uropathogens were frequently detected despite participants being recruited from the general population and not selected based on urinary symptoms or urological disease **(Table 2)**. *Enterococcus faecalis* and *Escherichia coli* were among the most prevalent uropathogens detected in the cohort, 95% and 48% of samples respectively, with similar prevalence among females and males (ca. 50%) **(Table 2)**. Other well-known uropathogens such as *Staphylococcus saprophyticus, Klebsiella pneumoniae, Pseudomonas aeruginosa* or *Proteus mirabilis* were not common in our cohort (≤ 15%) but they were more prevalent in samples from males **(Table 2)**.

**Table 2.** Prevalence and taxonomical classification of recognized uropathogens in the study population (1,615 participants). MGS = metagenomic species identifier; N = number of samples; % = percentage; *= former Gardnerella vaginalis.

| Total N | % | % females | Microorganism [Genus] [species] | Family | Order | Class | Phylum |
| --- | --- | --- | --- | --- | --- | --- | --- |
| 1537 | 95% | 54% | *Enterococcus faecalis* | Enterococcaceae | Lactobacillales | Bacilli | Bacillota |
| 1023 | 63% | 68% | *Streptococcus anginosus* | Streptococcaceae | Lactobacillales | Bacilli | Bacillota |
| 772 | 48% | 50% | *Escherichia coli* | Enterobacteriaceae | Enterobacterales | Gammaproteobacteria | Pseudomonadota |
| 562 | 35% | 65% | *Bifidobacterium vaginale* * | Bifidobacteriaceae | Actinomycetales | Actinomycetia | Actinomycetota |
| 456 | 28% | 65% | *Streptococcus agalactiae* | Streptococcaceae | Lactobacillales | Bacilli | Bacillota |
| 320 | 20% | 51% | *Staphylococcus aureus* | Staphylococcaceae | Staphylococcales | Bacilli | Bacillota |
| 240 | 15% | 41% | *Pseudomonas aeruginosa* | Pseudomonadaceae | Pseudomonadales | Gammaproteobacteria | Pseudomonadota |
| 212 | 13% | 37% | *Staphylococcus saprophyticus* | Staphylococcaceae | Staphylococcales | Bacilli | Bacillota |
| 198 | 12% | 36% | *Klebsiella pneumoniae* | Enterobacteriaceae | Enterobacterales | Gammaproteobacteria | Pseudomonadota |
| 77 | 5% | 54% | *Aerococcus urinae* | Aerococcaceae | Lactobacillales | Bacilli | Bacillota |
| 32 | 2% | 31% | *Proteus mirabilis* | Enterobacteriaceae | Enterobacterales | Gammaproteobacteria | Pseudomonadota |

Overall, we identified 3,668 microbial species belonging to 31 phyla, 48 classes, 109 orders, 256 families and 1,122 genera. Species assignment was achieved for 92% (3,367) of detected species, while 301 were assigned only to higher taxonomic ranks (275 to genus, 24 to family and 2 to order). Bacteria predominated across the cohort. Despite the large number of detected species, relatively few were consistently present across the population. A total of 26 species were detected in more than 50% of participants **(Table S2)**, whereas 415 species were detected in more than 5%, indicating that most detected species were uncommon. Among non-bacterial microorganisms, *Methanobrevibacter A smithi* was detected in 6% of participants (94% females) while fungi *Malassezia restricta* and *Malassezia globosa* were detected in 28% and 18% of participants, mostly males, whereas *Candida* species were rare (≈ 1% of participants).

Sex was the dominant determinant of urogenital microbial profiles whose differences could be seen across all taxonomical levels from phylum to species **(Figure 2)**. Female samples were predominantly characterized by *Lactobacillus* (*Lactobacillus iners, Lactobacillus crispatus, Lactobacilllus gasseri, Lactobacillus mulieris*), *Bifidobacterium (Bifidobacterium vaginale (*former *Gardnerella vaginalis*), *Bifidobacterium swidsinskii*, or *Bifidobacterium leopoldii), Streptococcus* (*Streptococcus anginosus, Streptococcus agalactiae*), whereas male samples showed greater heterogeneity and were predominantly characterized by Propionibacteriaceae, *Enterococcus faecalis, Corynebacterium falsenii, Neisseria suis*, and *Cutibacterium acnes*. A few samples were composed mainly by *Escherichia coli* in females while *Bifidobacterium swidwinskii, Bifidobacterium vaginale, Prevotella bivia, Streptococcus agalactiae*, or *Lactabacillus iners* were predominant bacteria in few male samples **(Figure 2)**.

**Figure 2:**
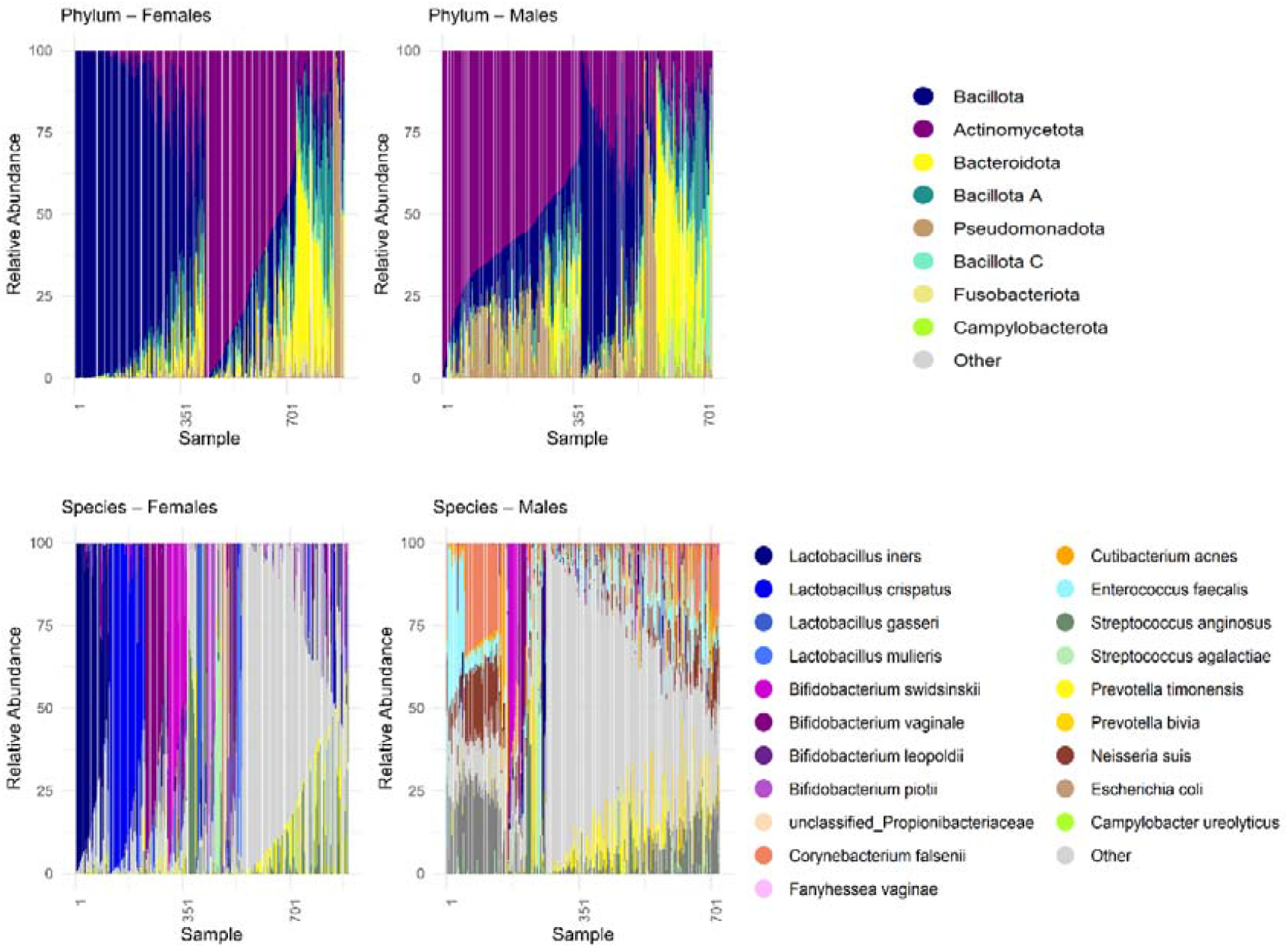
Microbial composition by sex and taxonomical levels phylum (top) and species (bottom). Stacked relative abundance of microorganism by sex and most abundant species in the sample. Samples are depicted in the horizontal (x) axis as lines and relative abundance is depicted in the vertical (y) axis.

### 3.3 Sex specific microbial profiles

Microbial diversity differed markedly by sex, whereas age showed only modest associations. Shannon diversity and inverse Simpson index were significantly higher in males after adjustment for age group and batch (Shannon diversity β = 0.51 (95% CI 0.42-0.60) and R^2^ = 0.08; inverse Simpson β = 1.23 (95% CI: 0.73-1.72) and R^2^ = 0.02; both p < 0.0001), whereas species richness did not differ by sex **(Figure 3)**. Participants aged 61-65 years had higher Shannon (β = 0.17 (95% CI 0.06-0.28), p = 0.003), and inverse Simpson diversity (β = 0.79 (95% CI 0.18-1.40), p = 0.011) compared to those aged 50-55 years and both older age groups showed higher richness (p < 0.05) independently of sex. Effect sizes for age were smaller than those observed for sex. In sensitivity analyses, results were consistent, but sex also remained significantly associated with richness.

**Figure 3:**
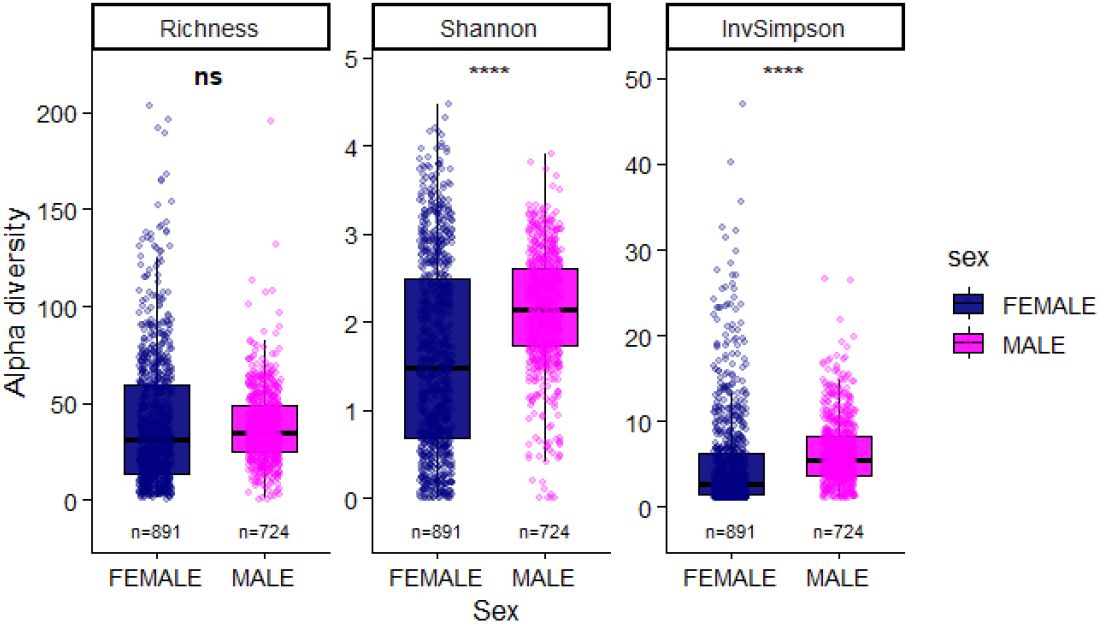
Alpha diversity indices in our cohort comparing females and males. (****) p-value < 0.0001

Sex explained substantially more variation than age across all beta-diversity metrics. Multivariate PERMANOVA adjusted for age and batch revealed statistically significant differences in beta diversity indices between sexes (Bray-Curtis R^2^ = 6.2%, Jaccard R^2^ = 5.2%, and Sørensen R^2^ = 8.5%; all p = 0.001) **(Figure 4)**. Homogeneity of dispersion was also significant with males having significantly lower dispersion than females; and thus, females had higher within-group variability (p < 0.001) suggesting that sex-associated differences reflect both shifts in community composition and differences in dispersion. Age was also statistically significant but explained a substantially smaller proportion of variation (<0.3% across metrics) while batch accounted for 2.5-5.7% of variation. Constrained ordination confirmed independent associations of sex, age, and batch with microbial community composition across all beta-diversity metrics (all p = 0.001), consistent with PERMANOVA results **(Figure S1)**. In sensitivity analyses including very low-biomass samples, the main effect of sex remained significant with similar effect sizes (Bray-Curtis R^2^ = 7.4%, Jaccard R^2^ = 6.0%, Sørensen R^2^ = 9.8%; all p = 0.001) and differences of dispersion between sexes also continued significant (p < 0.001). Overall, exclusion of very-low biomass samples did not materially alter the observed associations.

**Figure 4:**
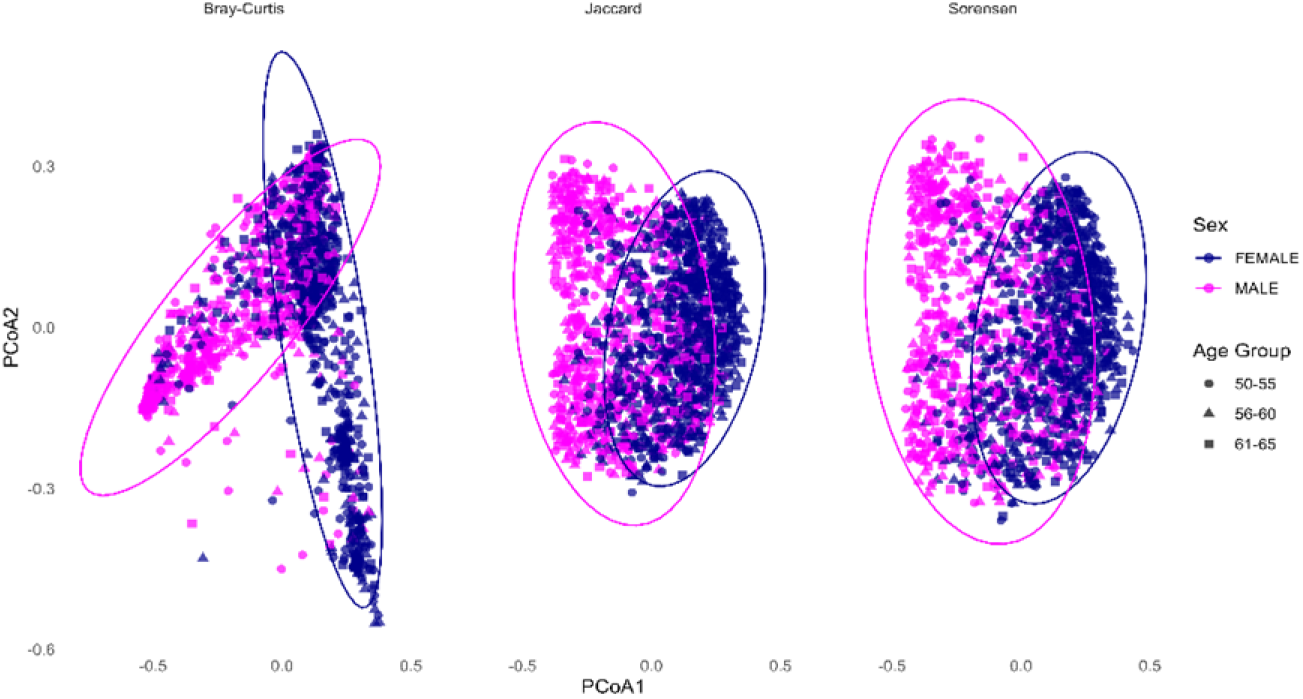
Principal coordinate analysis (PCoA) of beta diversity indices. Principal coordinate analyses based on Bray–Curtis (left), jaccard (middle), and Sørensen (right) dissimilarity matrices derived from multivariable PERMANOVA models adjusted for age group and batch. Points represent individual samples and are coloured by sex and shaped by age groups. Ellipses indicate 95% confidence intervals around group centroids.

For sex specific analyses, we applied a 5% prevalence exclusion filter as described in methods to avoid rare taxa emerging as false positives, which decreased our studied species to 648 species. Across multiple complementary statistical approaches, sex was consistently associated with marked differences in microbial composition.

Our indicator species analysis identified 437 species significantly associated with either sex (FDR adjusted p < 0.05). Of these, 67 species had an indicator value > 0.5, considered a strong association with a given group; and most were associated with females, and only eight with males **(Table S3)**. In our second exploratory analyses, LEfSe, we found 59 associations at different taxonomic levels differentially abundant in females and males **(Figure S2)** with genera such as *Lactobacillus, Bifidobacterium*, and *Escherichia* (including *Lactobacillus crispatus, Lactobacillus iners, Lactobacillus mulieris, Lactobacillus jensenni, Bifidobacterium vaginale* or *Escherichia coli*) enriched in females and *Citrobacter, Neisseria*, and *Alcalibacterium* (including *Citrobacter koseri* or *Neisseria suis*) enriched in males.

Differential abundance analyses consistently identified substantially more sex associated than age associated taxa. Sex and age showed minimal correlation (R = 0.06) and collinearity (VIF = 1.007), supporting variable independence and adjustment for age in differential abundance analyses. Across methods, substantially more taxa were associated with sex than age. DESeq2 identified 489 sex-associated species, decreasing to 392 after age adjustment, whereas age alone was associated with comparatively few taxa **(Figure S3** and **Figure S4)**.

ANCOM-BC, with strict Holm FDR, found 498 species associated with sexes, and it increased to 516 when including age adjustment. Only 78 species were independent of sex and significant in the age-only model. Finally, for sex tests ALDEx2 found 337 species differently abundant in either sex, while the model adjusting for age found 342 species. Eleven species were significantly associated with age; but only 5 remained significant in the age-only model and not in models that included sex. When comparing these methods with Spearman correlations, we observed that results were strongly correlated (DESeq2 and ANCOM-BC with rho = 0.91; followed by ANCOM-BC and ALDEx2 rho = 0.7; and by ALDEx2 and DESeq2 with rho = 0.66) **(Figure S5)**.

Using sex models adjusted for age, 66 species were consistent and significantly associated with females while 67 were for males across all differential abundance methods **(Tables S4** and **S5)**. Of these consistent species, 69 species showed a prevalence of 25% in either sex or in the whole cohort and are depicted in **Figure S6**, ranked by range and variance across the three analyses. A subset of 45 species is presented in **Figure 5**, displaying cohort prevalence and mean relative abundances for visualization.

**Figure 5:**
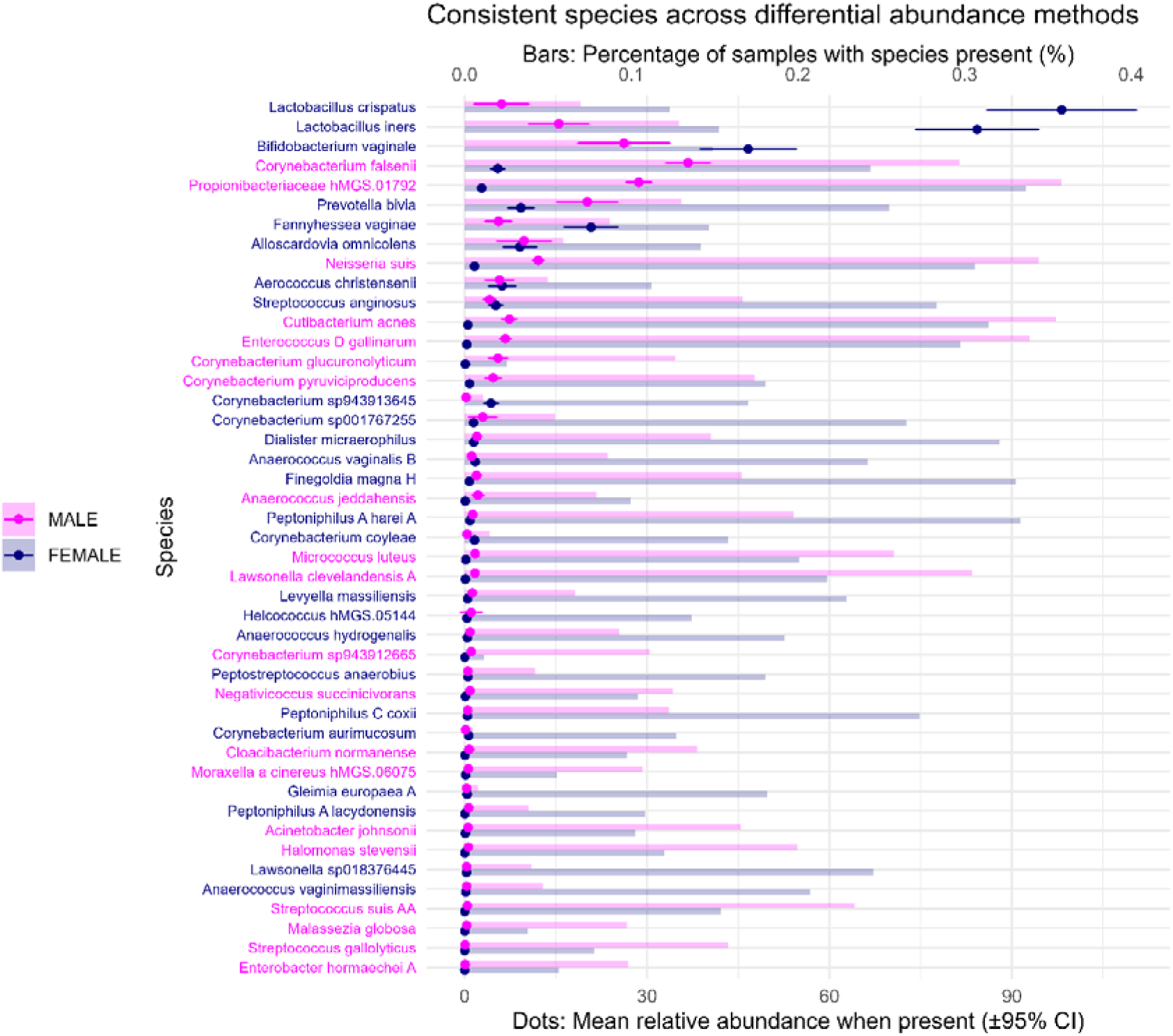
Species consistently identified across differential abundance methods. Bars represent species prevalence (percentage of samples in which the species was detected), and dots indicate mean relative abundance ±95% confidence intervals among samples in which the species was present. Species labels are coloured according to the sex in which the species exhibited higher abundance.

When adding to these three differential abundance methods the results of the indicator species and LefSe discriminant analyses, only 6 species were significantly consistent and all for females: *Aerococcus christensenii, Bifidobacterium vaginale, Fannyhessea vaginae* and three *Lactobacillus* (*L. iners, L. crispatus*, and *L. gasseri*). These 6 species consistent in all five methods suggest high-confidence specificity strengthening their role as potential key signatures for sex **(Table 4)**. *Lactobacillus mulieris* was significantly enriched in females across all methods except for ALDEx2 (Up in Male, effect 0.03).

**Table 4.**
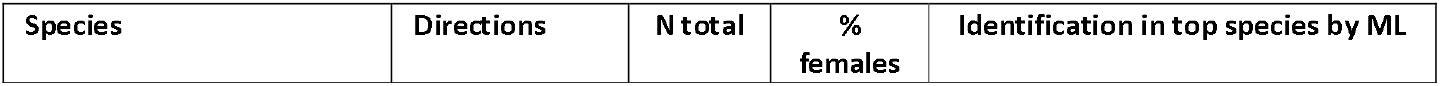
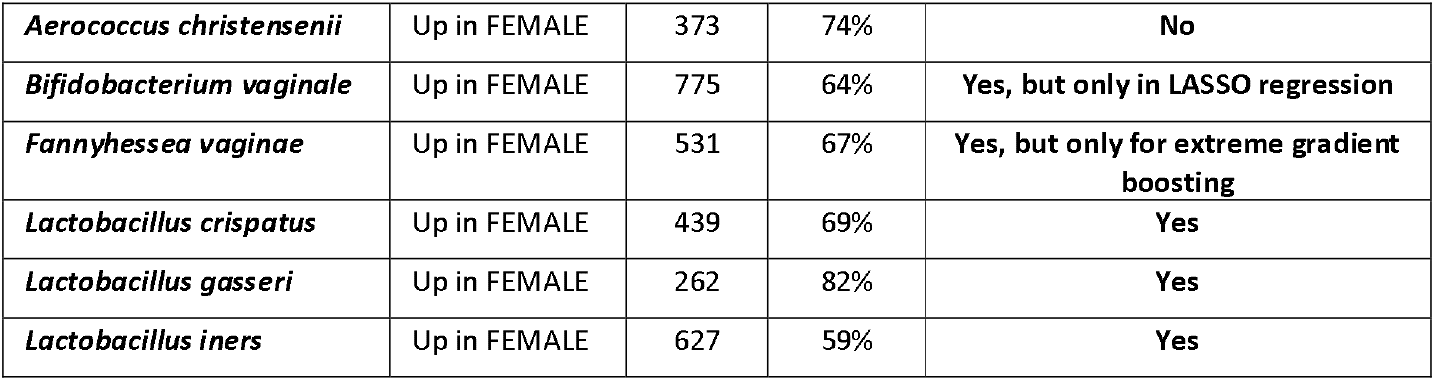
Microbial taxa associated with sex across all sex-specific analyses (indicator species, differential abundance, linear discriminant effect size analyses, ANCOM-BC and ALDEx2). MGS = Metagenomic species identifier, ML = supervised machine learning, N = number of samples, % = percentage

### 3.4 Machine learning validation of sex related microbial signatures

Three supervised machine learning models independently validated the observed sex specific microbial signatures from urine profiles with excellent discrimination (random forest AUC ≈ 0.99, extreme gradient boosting AUC ≈ 0.99, and LASSO regression AUC ≈ 0.95 **Figure S7)**. Performance dropped to chance level after label permutation (AUC dropped to ≈ 0.5; p value = 0.01), confirming that model performance was driven by sex rather than technical artefacts or random variation **(Figure S8)**. Fifteen taxa ranked among the top 50 predictors across all three models **(Figure 6)**. Of these 15, *Corynebacterium sp001767255, Lawsonella sp018376445, Gleimia europaea* A, *Dialister micraerophilus, UBA4285 sp900542465, Anaerococcus vaginalis* B, *Lactobacillus crispatus, Lactobacillus gassseri* and *Lactobacillus iners* were consistently identified and showed higher abundance in females in the LASSO model; whereas *Cutibacterium acnes, Enterococcus* D *gallinarum, Lawsonella clevelandensis* A, *Corynebacterium sp943912665, CAMCNU01* hMGS.01291, and *Propionimicrobium lymphophilum* showed higher abundance in males in the LASSO regression model. These findings closely matched the differential abundance analyses, further supporting robust sex-specific microbial signatures **(Figure 6** and **Table 4)**.

**Figure 6:**
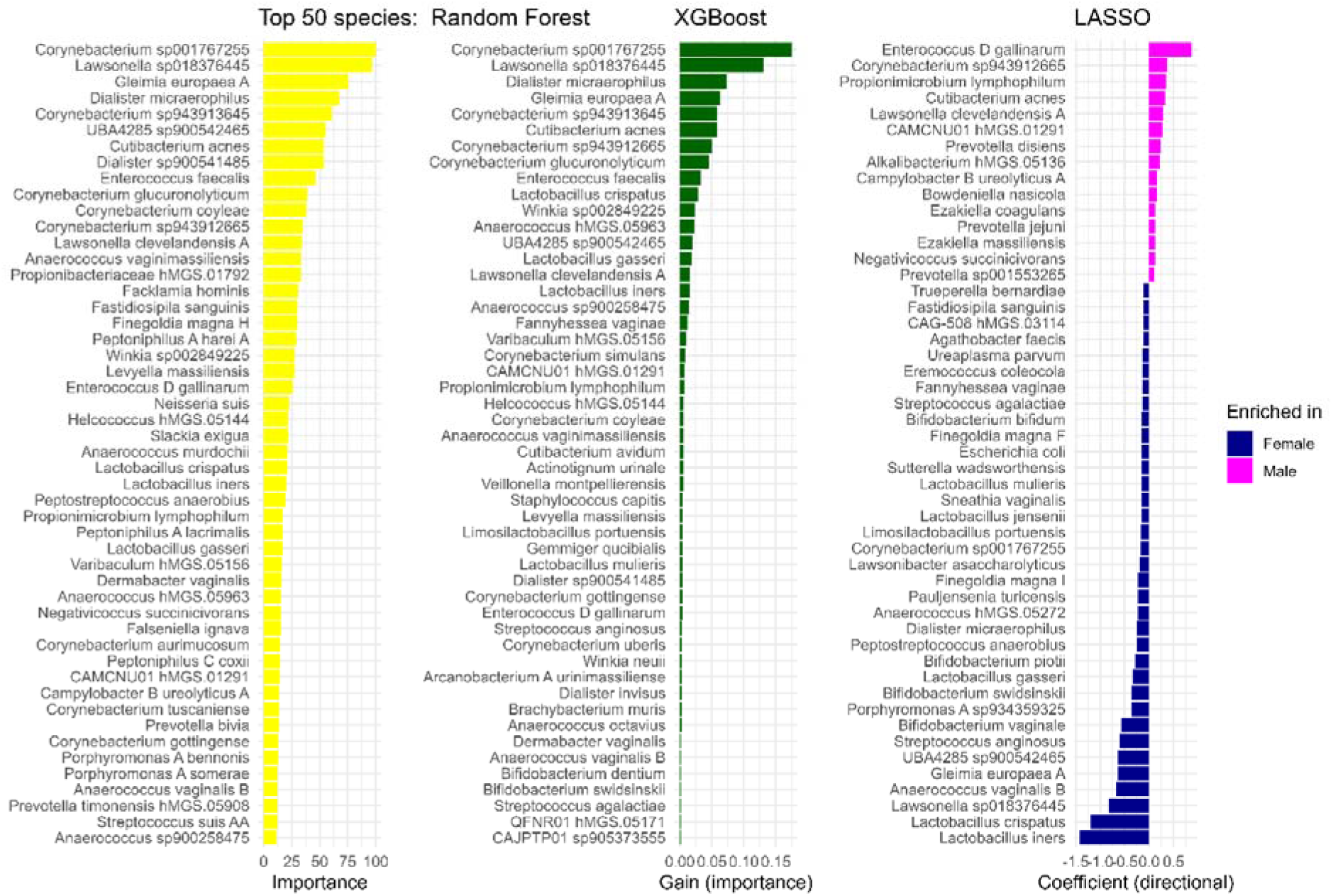
Top 50 taxa in each supervised machine learning approach. Random forest (importance in yellow); extreme gradient boosting (importance in green); and LASSO regression (blue for enrichment in females and magenta for males).

## 4. Discussion

This study investigated urine profiles and their microbial content in a large population-based study encompassing 1,615 SCAPIS participants. We provide the largest a population reference framework for microbial composition in voided urine in adults of 50-65 years without symptomatic UTIs [4]. As a result, five main findings emerge. First, recognized uropathogens are frequently detected in voided urine despite the absence of symptoms. Second, biological sex is the dominant determinant of microbial composition. Third, only a relatively small number of taxa consistently characterize urine profiles across the population despite the detection of thousands of microbial species. Fourth, lactobacilli and bifidobacteria predominated in female samples, suggesting the existence of a core microbiota alongside the high variability in low prevalence taxa; and fifth, supervised machine learning models were able to distinguish biological sex from urine profiles.

Notably, although participants were not selected based on urinary symptoms or urological disease, recognized uropathogens were frequently detected in urine. In some individuals, *Enterococcus faecalis, Escherichia coli*, or *Streptococcus anginosus* dominated urine microbial profiles and were among the most abundant species across the cohort. These observations suggest that the presence of recognized uropathogens in voided urine reflects microbial carriage rather than infection and reinforce that detection alone should not be interpreted as evidence of UTI in the absence of compatible symptoms or associated inflammation [24]. This is clinically relevant because UTIs remain among the most common bacterial infections worldwide, especially in peri- and post-menopausal women, accounting for a substantial proportion of antibiotic prescriptions and contributing to antimicrobial resistance [21,22]. However, the ecological distribution of recognized uropathogens varied between species, and for example, *Klebsiella pneumoniae*, an important reservoir of antimicrobial resistance [21], was more frequently detected in male urine profiles. Until recently, UTIs diagnosis relied primarily on urine culture, despite increasing evidence that standard culture captures a small fraction of microorganisms present in urine [13–15,23]. In addition, uropathogens may also be part of the vaginal and gut microbiome communities. These three biological niches share overlapping species with growing evidence linking their interactions to urological conditions [9,10,25]. Together, these findings support the need for population reference data when interpreting microorganisms detected in voided urine.

In our cohort, urine profiles from males exhibited higher Shannon and inverse Simpson diversity after adjustment for age and batch, whereas species richness did not differ by sex. Age was also independently associated with modest increases in diversity. Within the age range of the present study (50-65 years), sex explained substantially more microbial variation than age, which had comparatively modest effects (beta diversity metrics <0.3%). Biological sex was independently associated with microbial community composition, explaining approximately 5-8% of the variation in beta diversity metrics, with sex differences driven mainly by core taxa presence and abundance not by rare species. Sensitivity analyses including low-biomass samples showed comparable results for both alpha and beta diversity, indicating that exclusion of low-biomass samples did not materially alter the primary findings. Consequently, sex-related differences may be even more pronounced during the reproductive years, with more hormonal fluctuations, than observed in females around peri- and post-menopausal age. Future studies including younger women are needed to directly test this hypothesis.

Lactobacilli and bifidobacteria are part of the normal microbiota of the urinary tract, particularly in women [1] which is in line with our findings. Despite the use of different statistical approaches, 4 of the 6 species consistently significant for females belonged to these two groups. Lactobacilli were also of great importance when applying supervised machine learning approaches for differentiating females. *Lactobacillus iners* has been described as a transitional species in the vagina [26,27] but this was the species with highest average relative abundance in urine samples of peri- or menopausal females in our cohort. The second most abundant species in females, *Lactobacillus crispatus*, has been associated with vaginal and urinary health [27,28]. *Lactobacillus* abundance is associated with menopausal status and with reduced risk of urinary tract infection even though recent evidence suggest their potential role as pathobionts [29–32]. Our findings are also in accordance with previous studies reporting *Bifidobacterium* (a genus that includes the former *Gardnerella vaginalis*) being common “urotypes” in healthy females [29]. Approximately half of the female samples were dominated by a single species of *Lactobacillus* or *Bifidobacterium*. Other reported urotypes were dominated by *Streptococcus anginosus, Streptococcus agalactiae*, or *Escherichia coli*. Supervised machine learning pointed also to other bacteria important for female differentiation such as *Gleimia europaea* A or *Dialister micraerophilus*, both opportunistic pathogens associated with abscesses and UTIs in females [33,34].

Several species associated with males in our samples were previously linked to skin microbiota. This may partly reflect the low prevalence of circumcision in Sweden (approximately 5% of males) [35]. Hence, microorganisms could be easily transferred to urine from the penile oily or sebaceous skin, which is dominated by lipophilic microorganisms such as *Cutibacterium* spp. and *Malassezia* spp. [36]. Supervised machine learning models pointed to other skin microbes such as *Lawsonella clevelandensis*, an opportunistic pathogen associated with fatty soft skin abscesses [37]; *Cutibacterium acnes*, opportunistic pathogens associated with acne [38]; or *Propionimicrobium lymphophilum*, described to be part of the urinary tract and a beneficial commensal [24,39]. The latter study suggested that a more diverse microbiome is reflective of a urogenital healthy microbiome in males similarly to what has been seen in the gut [24]. These observations suggest that several taxa enriched in males may originate from skin, which may be consequence of the low percentage of circumcised males in Sweden, or represent commensal members of the male urogenital microbiota, although further work is needed to distinguish these possibilities.

Several highly prevalent taxa belonged to the Peptoniphilaceae family, members of which produce short-chain fatty acids with recognised roles in host-microbiota interactions mostly in the gut [40– 44]. Their potential contribution to urinary tract homeostasis including if and how these molecules are produced and act warrants further investigation.

### Strengths and limitations

To our knowledge, this is one of the largest and most comprehensive descriptions of microbial profiles in voided urine from a large population cohort free of urinary symptoms, which shows common uropathogens being detected in both female and male samples. First, studies published have mainly used 16S rRNA gene sequencing which often lacks resolution at the species level, while we reached species level in more than 90% of microorganisms. The use of shotgun sequencing allowed us to identify microorganisms in the kingdoms Archaea and Eukaryota even though most species belonged to Bacteria. Moreover, by including comparable numbers of females and males, we provide evidence of sex specific microbiome from urine profiles, which can be reliably distinguished using supervised machine-learning models. Collectively, these findings underscore the importance of incorporating sex as a key determinant in the design and interpretation of studies of urine samples and associated urinary/urogenital microbiome. To finish, voided urine samples offer practical advantages for clinical applications making them well suited for future research on urogenital microbiota signatures for a range of diseases.

Nonetheless, we also acknowledge some limitations. First, although our primary interest was the urinary microbiome, studying voided urine samples inevitably include microbial contributions from surrounding anatomical niches. Second, excluding very low biomass samples may have introduced selection bias, as these samples were predominantly from males. Hence, generalizability of microbial composition among males may be reduced although diversity associations were robust in sensitivity analyses. Third, our two batches differed in comorbidities, with participants in batch 2 exhibiting a higher prevalence of diabetes, prior cardiovascular disease, and greater coronary artery calcium severity. Although all primary models were adjusted for batch, residual confounding related to comorbidity clustering cannot be entirely excluded. Importantly, batch explained less variation in microbial composition than sex, indicating that sex-associated differences were not primarily attributable to batch effects. Lastly, taxonomical annotation of bacteria by CHAMP used the release 214 of the GTDB database, and for which a 2026 update will be shortly released [18,45]. Some species were novel relative to GTDB and therefore lack official taxonomic names and they remain poorly characterized; in such cases, MGS identifiers must be used, which prevent from having firm conclusions.

Investigating additional factors that may influence the composition and function of the urinary or urogenital microbiome was beyond the scope of this study, which focused on characterizing urine microbiome profiles in a Swedish population without urinary symptoms. Moreover, the mechanisms by which other factors shape the urinary microbiome are not yet fully understood and such factors warrant separate investigation (e.g., hypertension, kidney damage, or glycosuria) [46,47].

## 5. Conclusions

This study provides a population reference for microbial profiles in voided urine and substantially advances current understanding of the urogenital microbiome. We demonstrate that recognized uropathogens are frequently detected in individuals without symptomatic UTIs. Our findings suggest that the urogenital microbiota is more diverse than previously assumed and that exhibits important sex differences. Biological sex emerged as the principal determinant of microbial composition, with lactobacilli and bifidobacteria predominating in female urine profiles. Finally, we highlight the importance in incorporating biological sex in future studies to better understand how urine microbiome profiles vary in the context of diseases and clarify its role in health and disease.

## 6. Declarations

### Author Contributions

Conceptualization TG, JÄ; microbiome sequencing and data annotation OL, PRG, JMM; data analysis TG; writing original draft TG; writing—review and editing TG, YTL, TS, SA, TF, JÄ, JMM, HBN. All authors have read and agreed to the published version of the manuscript.

## Acknowledgements

Funding: We acknowledge funding from the Swedish research council (Vetenskapsrådet grant number 2019-01015 (J.Ä.), 2019-01471 (T.F.) and 2020-0243 (J.Ä.)), from the Swedish heart lung foundation (Hjärt-Lungfonden grant number 2021-0357 (J.Ä.), 2023-0687 (T.F.) and 2024-0486 (J.Ä.)), and from Center of clinical research (CKF) in Region Dalarna, Falun, Sweden. Dr Ahmad was supported by research grants from FORMAS–Early Career Grant (no. 2020-00989), Swedish Research Council-Early Career Grant (no. 2022-01460), EFSD/Novo Nordisk and EpiHealth. SCAPIS has been primarily funded by the Swedish Heart-Lung Foundation (Hjärt-Lungfonden), and other funders include the Knut and Alice Wallenberg Foundation, the Swedish Research Council, and VINNOVA. SCAPIS also acknowledges the participation of over 30,000 volunteers, as well as the collaboration of the six host universities and university hospitals: University of Gothenburg and Sahlgrenska University Hospital; Karolinska Institutet and Karolinska University Hospital; Linköping University and Linköping University Hospital; Lund University and Skåne University Hospital; Umeå University and Umeå University Hospital; Uppsala University and Uppsala University Hospital. This project is part of SCAPIS petition 210 and acknowledges the contribution of the Uppsala node. The funders had no role in study design, data collection and analysis, decision to publish, or preparation of the manuscript.

## Ethics approval and consent to participate

SCAPIS was approved by the ethics committee at Umeå University (DNR 2010-228-31M), and it was adhered to the Declaration of Helsinki. The present study was approved by the Swedish Ethical Review Authority (DNR 2018-315; SCAPIS petition 210). All participants provided written informed consent.

## Consent for publication

All participants provided written informed consent.

## Availability of data

Authors Tiscar Graells and Johan Ärnlöv had full access to all the data in the study and takes responsibility for the integrity of the data and the accuracy of the data analysis. The data underlying this study are derived from the Swedish CArdioPulmonary bioImage Study (SCAPIS) and include human data classified as sensitive personal data under the EU General Data Protection Regulation (GDPR) and Swedish law. As such, individual-level data cannot be made publicly available. Data are available through a controlled access process managed by the SCAPIS steering committee. Qualified researchers may apply for access following approval by the Swedish Ethical Review Authority and the SCAPIS data access committee. Information on the application procedure is available at the SCAPIS data access board (https://www.scapis.org/how-to-apply/data-access-board/).

