## Supplementary Information for "Marked sex differences and frequent carriage of recognized uropathogens in the human urogenital microbiome: urine profiles from a population-level shotgun metagenomics study"

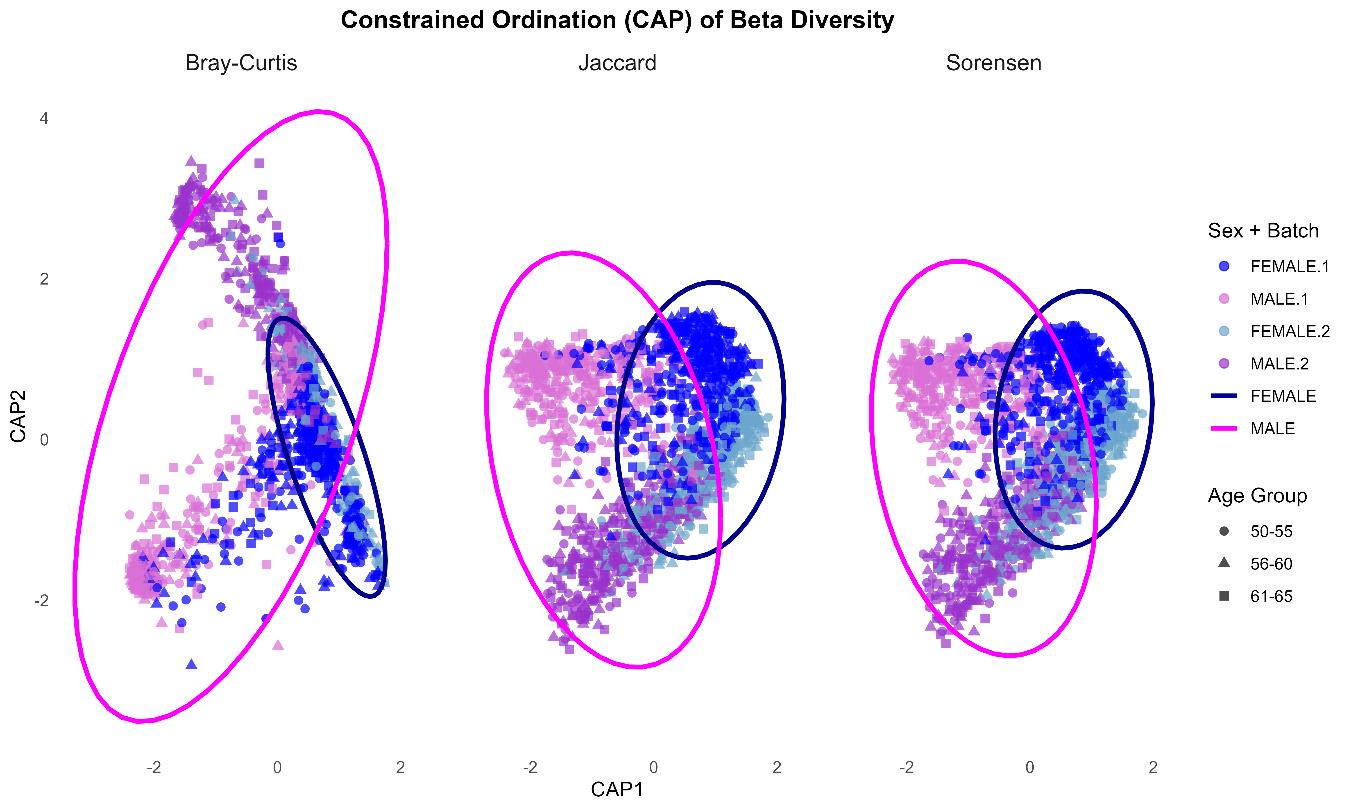

**Figure S1:** Constrained ordination of principal coordinates (CAP) based on Bray–Curtis (left), Jaccard (middle), and Sørensen (right) dissimilarity matrices from models including sex, age group, and batch. Points represent individual samples and are coloured by sex-batch combinations and are shaped by age group. Ellipses indicate 95% confidence intervals around group centroids for sex.

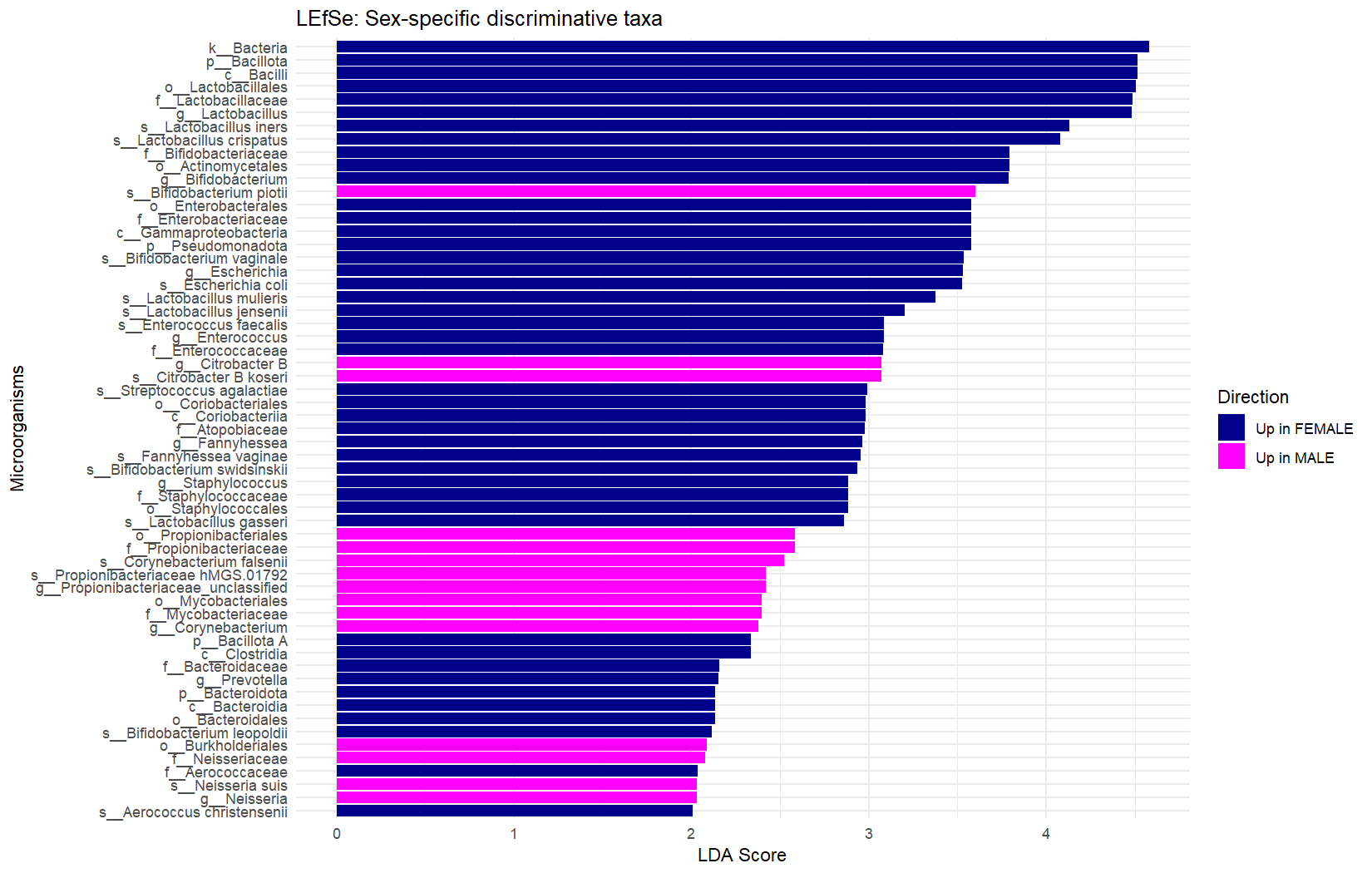

**Figure S2:** Bar plot of linear discriminant different abundant (LefSE) ordered by their score (LDA > 2) and q-value (FDR adjusted p-value).

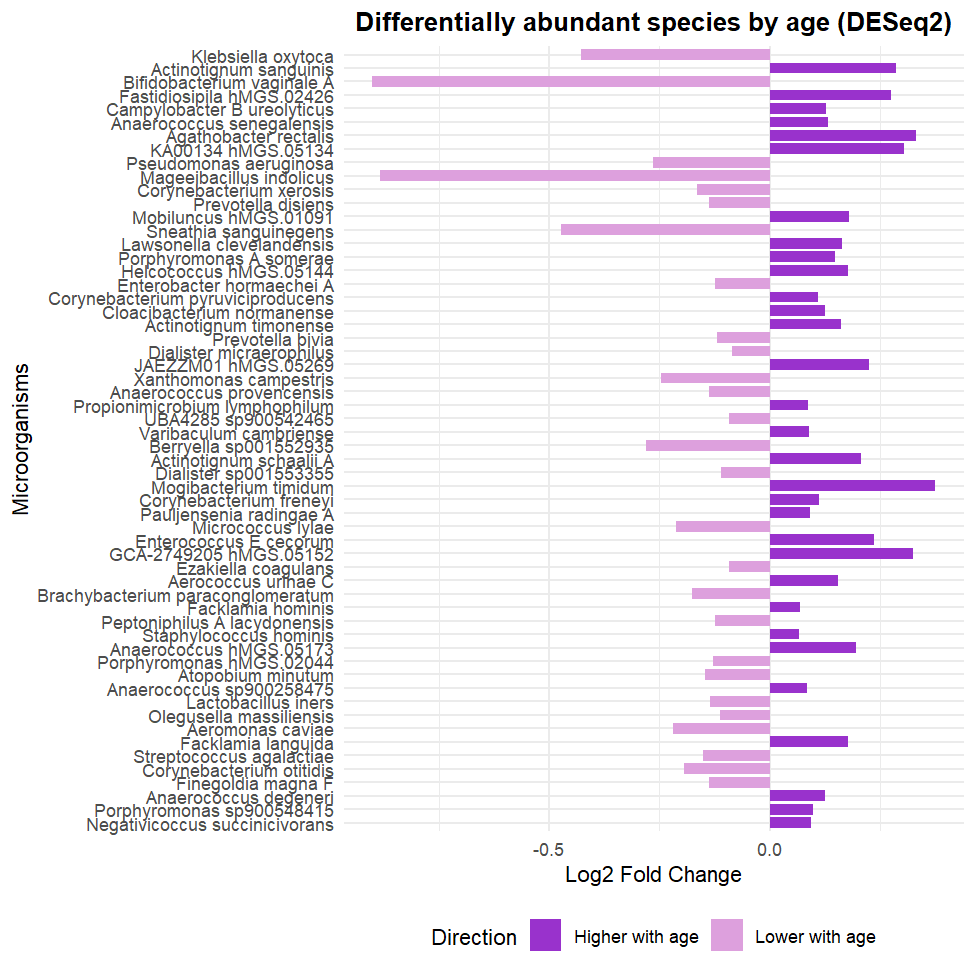

**Figure S3:** Bar plot of the differentially abundant species (DeSEq2) by age (batch adjusted) and ordered by ascending q-value (FDR adjusted p-value).

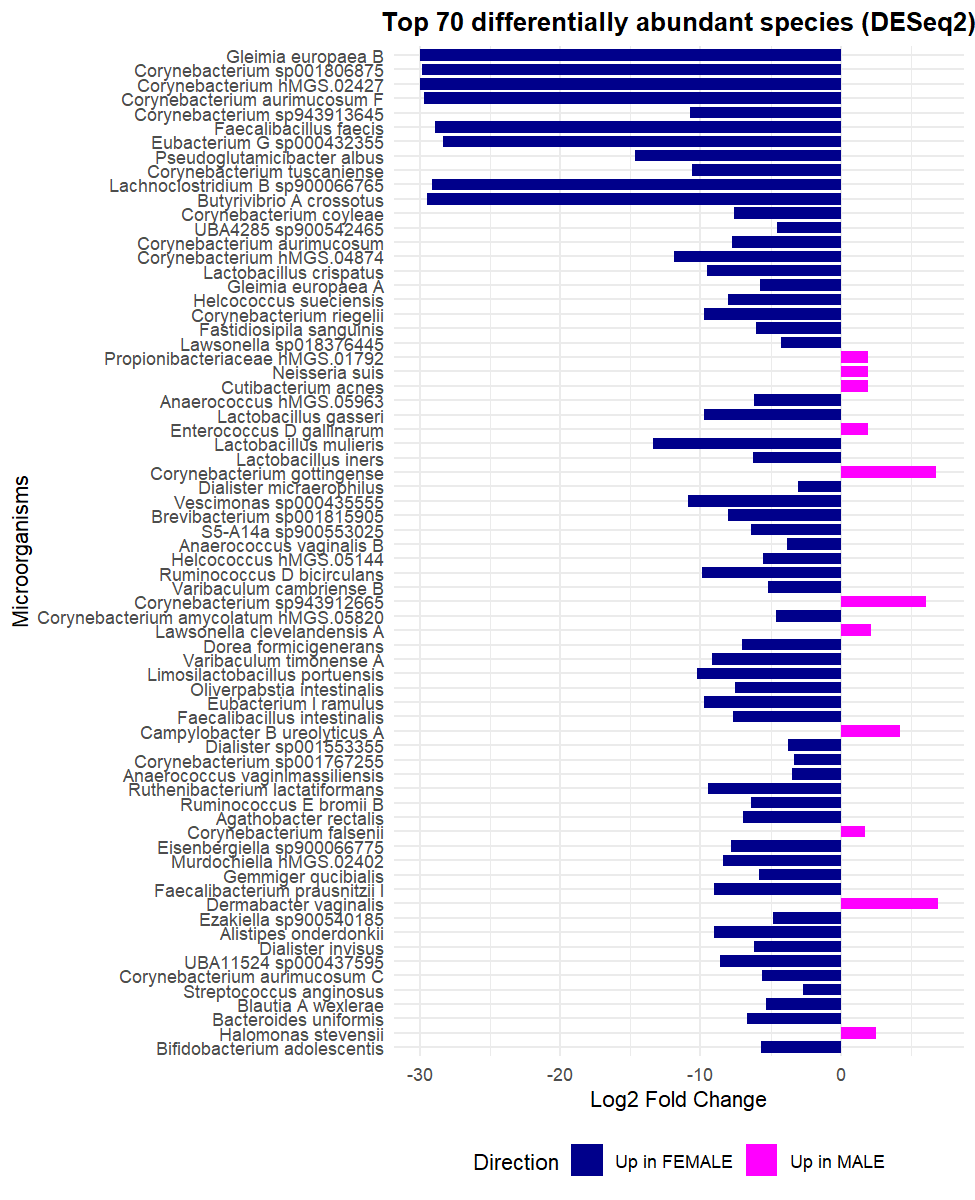

**Figure S4:** Bar plot of the top 70 differentially abundant microorganisms (DeSEq2) by sex (batch and age adjusted) and ordered by ascending q-value (FDR adjusted p-value).

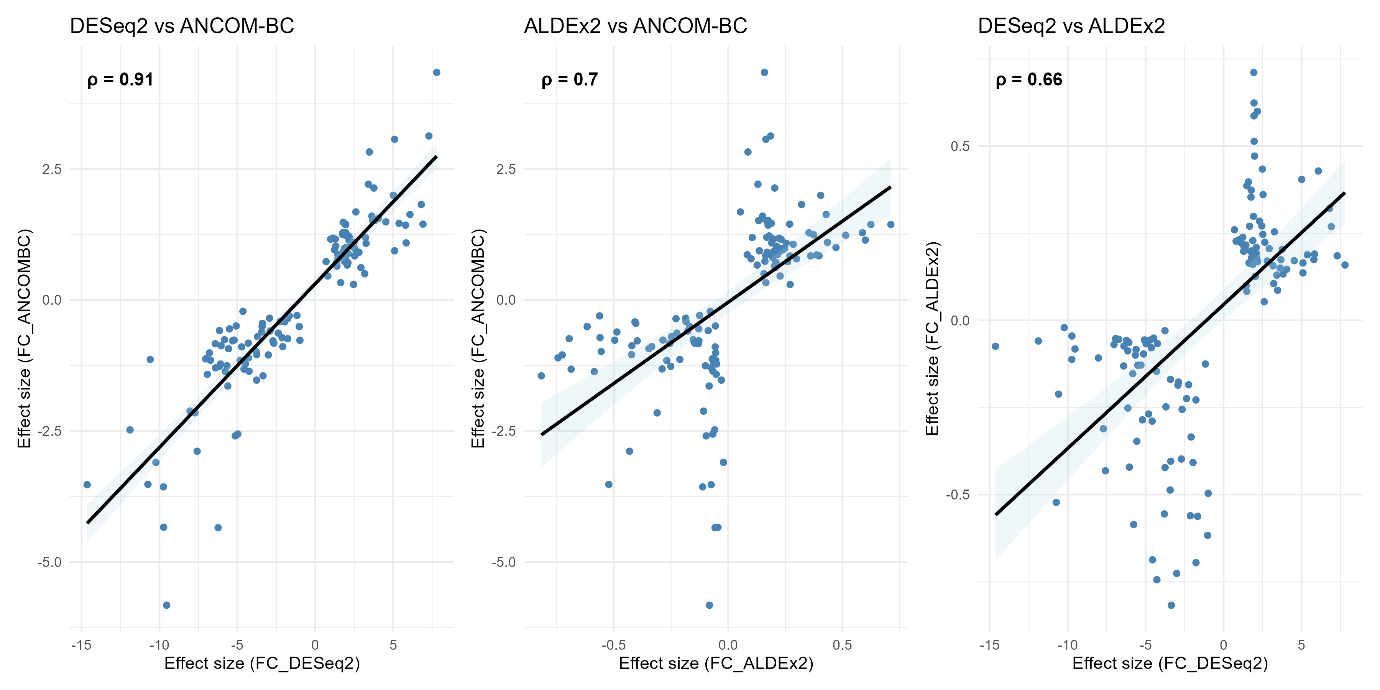

**Figure S5:** Pairwise spearman correlations between the three differential abundance methods tested: DESeq2, ANCOM-BC and ALDEx2

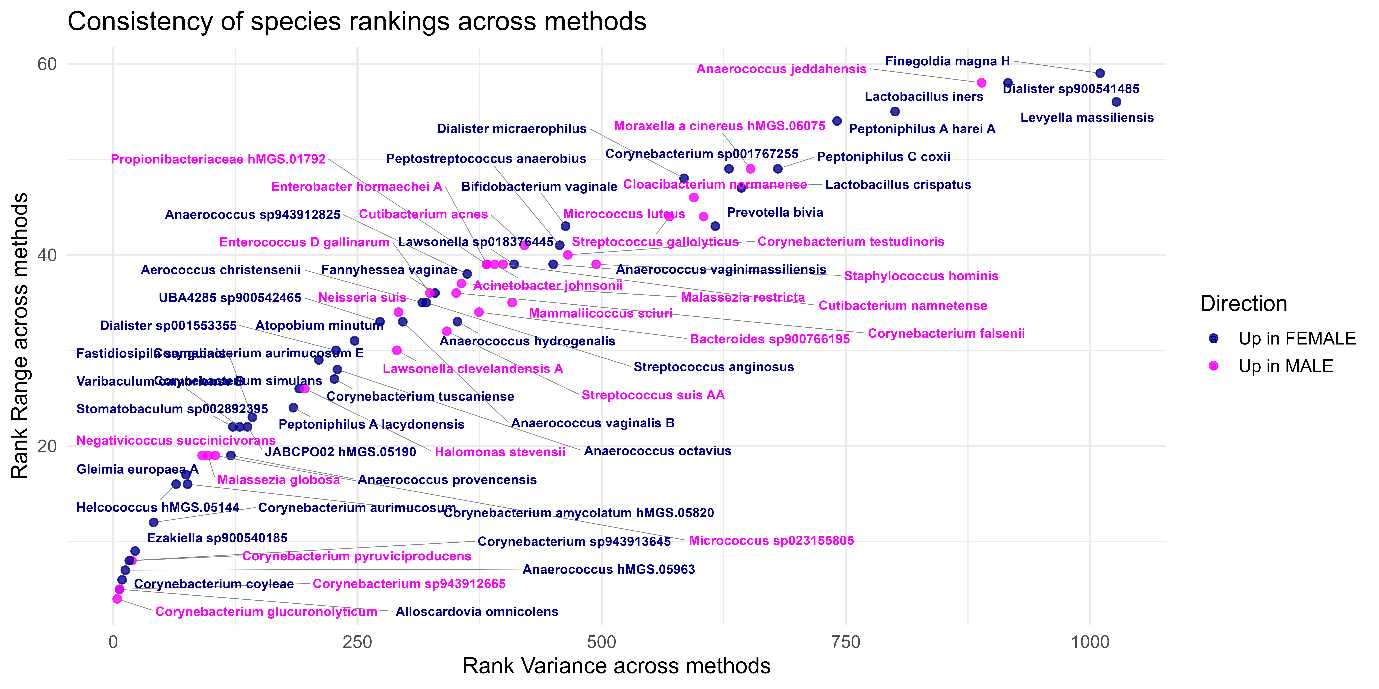

**Figure S6**: Scatter plot of species associated with either sex according to ranks in range and variance across the three sex-specific differential abundance analyses adjusted for age (differential abundance DeSeq2, ANCOM-BC and ALDEx2). We have only pictured species present in more than 25% of samples in the whole cohort or either sex. Species placed on the left and the bottom quarter (low variance and low range) are the most consistently ranked across DESeq2, ALDEx2, and ANCOM-BC.

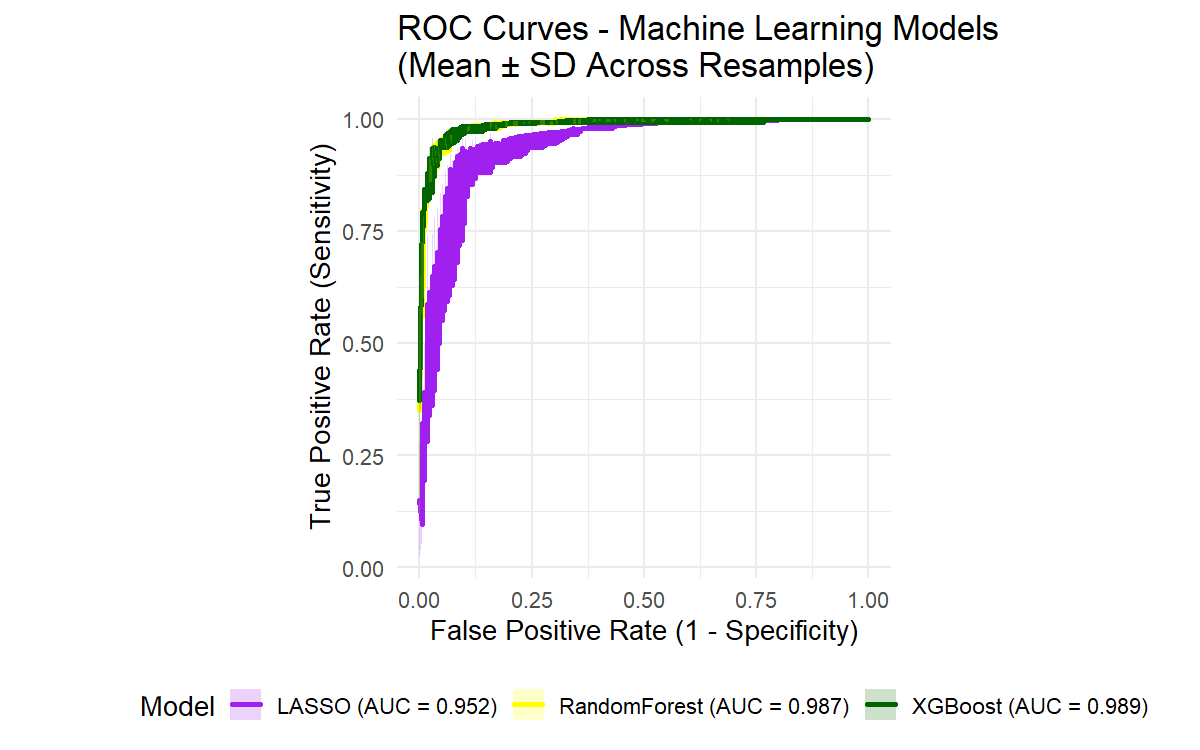
**Figure S7:** Area under the ROC curves (AUC) depicting supervised machine learning model performances, sensitivity, and specificity.

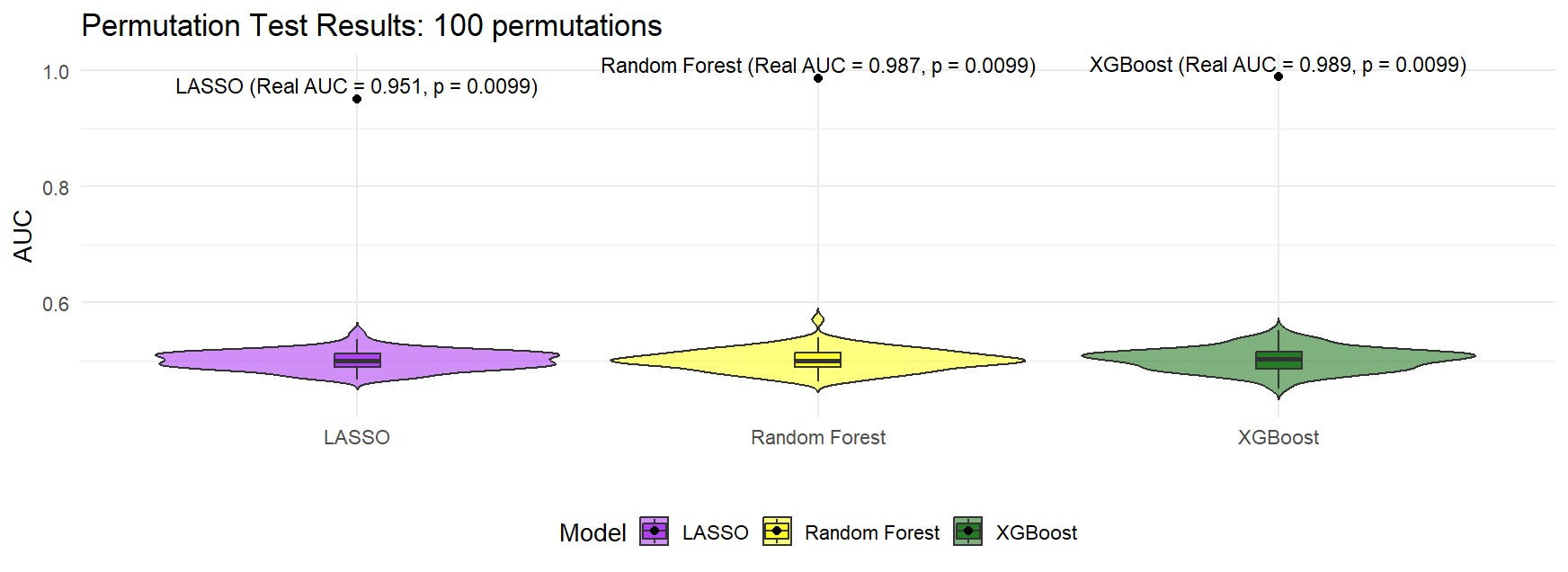

**Figure S8:** Permutation test results with 100 permutations shuffling labels. AUC values of the 100 permutations are depicted in violin plots, the average AUC is depicted by the line in boxplots and real AUC is depicted by the black dot (all outside the violin plots).

**Supplementary Tables**

**Table S1:** Distribution of major comorbidities by batch in our cohort.

|  | ALL (N = 1615) | BATCH 1 (N=800) | BATCH 2 (N=815) |
| --- | --- | --- | --- |
| Diabetes | 204 (12.6%) | 65 (8.1%) | 139 (17.1%) |
| Chronic kidney disease |  |  |  |
| None or stage 1 | 760 (47.1%) | 357 (44.6%) | 403 (49.4%) |
| Stage 2, 3 or 4 | 854 (52.9%) | 443 (55.4%) | 411 (50.4%) |
| Previous cardiovascular events (questionnaire) | 141 (8.7%) | 40 (5%) | 101 (12.4%) |
| Subclinical atherosclerosis  (by Coronary Artery Calcium score) |  |  |  |
| None | 947 (58.6%) | 515 (64.4%) | 432 (53%) |
| Mild | 403 (25%) | 188 (23.5%) | 215 (26.4%) |
| Moderate | 151 (9.3%) | 60 (7.5%) | 91 (11.2%) |
| Severe | 114 (7%) | 37 (4.6%) | 77 (9.4%) |

**Table S2:** Taxonomical classification of the 26 species that were present in more than half of the samples. MGS = metagenomic species identifier; N = number of samples, % = percentage

| **N** | **%**  **females** | **MGS** | **Species** | **Genus** | **Family** | **Order** | **Class** | **Phylum** |
| --- | --- | --- | --- | --- | --- | --- | --- | --- |
| 1537 | 54% | hMGS.00092 | ***Enterococcus faecalis*** | *Enterococcus* | Enterococcaceae | Lactobacillales | Bacilli | Bacillota |
| 1533 | 54% | hMGS.01792 | **Propionibacteriaceae hMGS.01792** | *unclassified* | Propionibacteriaceae | Propionibacteriales | Actinomycetia | Actinomycetota |
| 1472 | 52% | hMGS.00201 | ***Cutibacterium acnes*** | *Cutibacterium* | Propionibacteriaceae | Propionibacteriales | Actinomycetia | Actinomycetota |
| 1431 | 52% | hMGS.00652 | ***Neisseria suis*** | *Neisseria* | Neisseriaceae | Burkholderiales | Gammaproteobacteria | Pseudomonadota |
| 1399 | 52% | hMGS.00682 | ***Enterococcus D gallinarum*** | *Enterococcus D* | Enterococcaceae | Lactobacillales | Bacilli | Bacillota |
| 1218 | 67% | hMGS.05908 | ***Prevotella timonensis* hMGS.05908** | *Prevotella* | Bacteroidaceae | Bacteroidales | Bacteroidia | Bacteroidota |
| 1206 | 67% | hMGS.00790 | ***Peptoniphilus A harei A*** | *Peptoniphilus A* | Peptoniphilaceae | Tissierellales | Clostridia | Bacillota A |
| 1184 | 50% | hMGS.03746 | ***Corynebacterium falsenii*** | *Corynebacterium* | Mycobacteriaceae | Mycobacteriales | Actinomycetia | Actinomycetota |
| 1138 | 72% | hMGS.00817 | ***Dialister sp900541485*** | *Dialister* | Dialisteraceae | Veillonellales | Negativicutes | Bacillota C |
| 1138 | 71% | hMGS.01330 | ***Finegoldia magna H*** | *Finegoldia* | Peptoniphilaceae | Tissierellales | Clostridia | Bacillota A |
| 1135 | 47% | hMGS.04022 | ***Lawsonella clevelandensis A*** | *Lawsonella* | Mycobacteriaceae | Mycobacteriales | Actinomycetia | Actinomycetota |
| 1125 | 67% | hMGS.00758 | ***Winkia sp002849225*** | *Winkia* | Actinomycetaceae | Actinomycetales | Actinomycetia | Actinomycetota |
| 1091 | 58% | hMGS.00084 | ***Staphylococcus epidermidis*** | *Staphylococcus* | Staphylococcaceae | Staphylococcales | Bacilli | Bacillota |
| 1077 | 73% | hMGS.00576 | ***Dialister micraerophilus*** | *Dialister* | Dialisteraceae | Veillonellales | Negativicutes | Bacillota C |
| 1023 | 68% | hMGS.00468 | ***Streptococcus anginosus*** | *Streptococcus* | Streptococcaceae | Lactobacillales | Bacilli | Bacillota |
| 1003 | 62% | hMGS.00350 | ***Propionimicrobium lymphophilum*** | *Propionimicrobium* | Propionibacteriaceae | Propionibacteriales | Actinomycetia | Actinomycetota |
| 1001 | 49% | hMGS.00154 | ***Micrococcus luteus*** | *Micrococcus* | Micrococcaceae | Actinomycetales | Actinomycetia | Actinomycetota |
| 992 | 74% | hMGS.01897 | ***Facklamia hominis*** | *Facklamia* | Aerococcaceae | Lactobacillales | Bacilli | Bacillota |
| 985 | 73% | hMGS.00892 | ***Peptoniphilus A lacrimalis*** | *Peptoniphilus A* | Peptoniphilaceae | Tissierellales | Clostridia | Bacillota A |
| 925 | 69% | hMGS.00574 | ***Campylobacter B ureolyticus*** | *Campylobacter B* | Campylobacteraceae | Campylobacterales | Campylobacteria | Campylobacterota |
| 910 | 73% | hMGS.01075 | ***Peptoniphilus C coxii*** | *Peptoniphilus C* | Peptoniphilaceae | Tissierellales | Clostridia | Bacillota A |
| 907 | 65% | hMGS.01637 | ***Anaerococcus senegalensis*** | *Anaerococcus* | Peptoniphilaceae | Tissierellales | Clostridia | Bacillota A |
| 905 | 76% | hMGS.02977 | ***UBA4285 sp900542465*** | *UBA4285* | Lachnospiraceae | Lachnospirales | Clostridia | Bacillota A |
| 882 | 52% | hMGS.00145 | ***Staphylococcus hominis*** | *Staphylococcus* | Staphylococcaceae | Staphylococcales | Bacilli | Bacillota |
| 880 | 71% | hMGS.00191 | ***Prevotella bivia*** | *Prevotella* | Bacteroidaceae | Bacteroidales | Bacteroidia | Bacteroidota |
| 839 | 45% | hMGS.05935 | ***Streptococcus suis AA*** | *Streptococcus* | Streptococcaceae | Lactobacillales | Bacilli | Bacillota |

**Table S3:** Strong indicator taxa for each sex resulting from the indicator specific analysis (indicator value (stat) > 0.5, index = 1 (indicator for females) and index =2 (indicator for males)). MGS = metagenomic species identifier. All p.values adjusted by false discovery rate were ≤ 0.001.

| **Species** | **MGS** | **s.Female** | **s.Male** | **Index** | **stat** |
| --- | --- | --- | --- | --- | --- |
| UBA4285 sp900542465 | hMGS.02977 | 1 | 0 | 1 | 0.85 |
| Dialister micraerophilus | hMGS.00576 | 1 | 0 | 1 | 0.85 |
| Dialister sp900541485 | hMGS.00817 | 1 | 0 | 1 | 0.84 |
| Streptococcus anginosus | hMGS.00468 | 1 | 0 | 1 | 0.81 |
| Corynebacterium sp001767255 | hMGS.03760 | 1 | 0 | 1 | 0.8 |
| Lawsonella sp018376445 | hMGS.01923 | 1 | 0 | 1 | 0.79 |
| Anaerococcus vaginalis B | hMGS.00945 | 1 | 0 | 1 | 0.79 |
| Peptoniphilus C coxii | hMGS.01075 | 1 | 0 | 1 | 0.75 |
| Anaerococcus vaginimassiliensis | hMGS.03324 | 1 | 0 | 1 | 0.74 |
| Streptococcus suis AA | hMGS.05935 | 0 | 1 | 2 | 0.72 |
| Atopobium deltae | hMGS.00904 | 1 | 0 | 1 | 0.72 |
| Prevotella bergensis | hMGS.07662 | 1 | 0 | 1 | 0.7 |
| Anaerococcus murdochii | hMGS.06888 | 1 | 0 | 1 | 0.7 |
| Gleimia europaea A | hMGS.01062 | 1 | 0 | 1 | 0.69 |
| Dialister sp001553355 | hMGS.00496 | 1 | 0 | 1 | 0.69 |
| Halomonas stevensii | hMGS.06317 | 0 | 1 | 2 | 0.69 |
| Pauljensenia turicensis | hMGS.00437 | 1 | 0 | 1 | 0.69 |
| Corynebacterium sp943913645 | hMGS.05841 | 1 | 0 | 1 | 0.68 |
| Slackia exigua | hMGS.01595 | 1 | 0 | 1 | 0.68 |
| Mobiluncus hMGS.01090 | hMGS.01090 | 1 | 0 | 1 | 0.68 |
| Peptostreptococcus anaerobius | hMGS.00580 | 1 | 0 | 1 | 0.67 |
| Varibaculum massiliense | hMGS.01220 | 1 | 0 | 1 | 0.67 |
| Levyella massiliensis | hMGS.00726 | 1 | 0 | 1 | 0.67 |
| Corynebacterium aurimucosum E | hMGS.01113 | 1 | 0 | 1 | 0.66 |
| Corynebacterium simulans | hMGS.01240 | 1 | 0 | 1 | 0.66 |
| Corynebacterium coyleae | hMGS.01419 | 1 | 0 | 1 | 0.66 |
| Prevotella buccalis | hMGS.00976 | 1 | 0 | 1 | 0.66 |
| Lactobacillus iners | hMGS.00136 | 1 | 0 | 1 | 0.64 |
| Fastidiosipila sanguinis | hMGS.07481 | 1 | 0 | 1 | 0.64 |
| Acinetobacter johnsonii | hMGS.01295 | 0 | 1 | 2 | 0.63 |
| Porphyromonas A sp001808555 | hMGS.02291 | 1 | 0 | 1 | 0.63 |
| Anaerococcus sp943912825 | hMGS.07406 | 1 | 0 | 1 | 0.62 |
| Citrobacter B koseri | hMGS.00719 | 0 | 1 | 2 | 0.62 |
| Corynebacterium amycolatum | hMGS.05820 | 1 | 0 | 1 | 0.61 |
| Fannyhessea vaginae | hMGS.00196 | 1 | 0 | 1 | 0.61 |
| Porphyromonas A bennonis | hMGS.01359 | 1 | 0 | 1 | 0.61 |
| Bifidobacterium vaginale | hMGS.00248 | 1 | 0 | 1 | 0.61 |
| Alloscardovia omnicolens | hMGS.00407 | 1 | 0 | 1 | 0.61 |
| Peptoniphilus E obesi | hMGS.04220 | 1 | 0 | 1 | 0.61 |
| Helcococcus hMGS.05144 | hMGS.05144 | 1 | 0 | 1 | 0.61 |
| Corynebacterium aurimucosum | hMGS.03743 | 1 | 0 | 1 | 0.59 |
| Peptoniphilus A grossensis | hMGS.01356 | 1 | 0 | 1 | 0.59 |
| Corynebacterium glucuronolyticum | hMGS.01318 | 0 | 1 | 2 | 0.58 |
| Anaerococcus hMGS.05963 | hMGS.05963 | 1 | 0 | 1 | 0.58 |
| Lactobacillus crispatus | hMGS.00239 | 1 | 0 | 1 | 0.58 |
| Winkia neuii | hMGS.05743 | 1 | 0 | 1 | 0.57 |
| Ezakiella sp900540185 | hMGS.01327 | 1 | 0 | 1 | 0.56 |
| Streptococcus gallolyticus | hMGS.00937 | 0 | 1 | 2 | 0.56 |
| Streptococcus agalactiae | hMGS.00714 | 1 | 0 | 1 | 0.56 |
| Malassezia restricta | hMGS.10057 | 0 | 1 | 2 | 0.56 |
| Corynebacterium tuscaniense | hMGS.01241 | 1 | 0 | 1 | 0.55 |
| Peptococcus niger | hMGS.01355 | 1 | 0 | 1 | 0.55 |
| Falseniella ignava | hMGS.06509 | 1 | 0 | 1 | 0.55 |
| Corynebacterium sp943912665 | hMGS.07606 | 0 | 1 | 2 | 0.55 |
| Atopobium minutum | hMGS.02095 | 1 | 0 | 1 | 0.55 |
| Olegusella massiliensis | hMGS.02268 | 1 | 0 | 1 | 0.54 |
| Stomatobaculum sp002892395 | hMGS.00834 | 1 | 0 | 1 | 0.54 |
| Varibaculum cambriense B | hMGS.03234 | 1 | 0 | 1 | 0.54 |
| Anaerococcus octavius | hMGS.01500 | 1 | 0 | 1 | 0.53 |
| Aerococcus christensenii | hMGS.00589 | 1 | 0 | 1 | 0.53 |
| Anaerococcus provencensis | hMGS.03321 | 1 | 0 | 1 | 0.53 |
| Porphyromonas hMGS.02044 | hMGS.02044 | 1 | 0 | 1 | 0.52 |
| JABCPO02 | hMGS.05190 | 1 | 0 | 1 | 0.52 |
| Berryella sp900604965 | hMGS.02101 | 1 | 0 | 1 | 0.52 |
| Helcococcus sueciensis | hMGS.07846 | 1 | 0 | 1 | 0.52 |
| S5-A14a sp000758905 | hMGS.04351 | 1 | 0 | 1 | 0.51 |
| CAMCNU01 hMGS.05310 | hMGS.05310 | 1 | 0 | 1 | 0.5 |

**Table S4:** Agreement between the three differential abundance methods (DESeq2, ANCOMBC, ALDEx2) of species consistent and significantly associated with females (higher abundance in females). FC = Fold change

| **Species** | **MGS** | **FC_DESeq2** | **FC_ANCOMBC** | **FC_ALDEx2** |
| --- | --- | --- | --- | --- |
| Agathobacter rectalis | hMGS.00001 | -6,930375587 | -1,419060378 | -0,052290264 |
| Bifidobacterium longum | hMGS.00002 | -4,724456358 | -0,895288349 | -0,056796463 |
| Gemmiger qucibialis | hMGS.00003 | -5,830018133 | -0,758770307 | -0,152986746 |
| Bifidobacterium adolescentis | hMGS.00004 | -5,662413142 | -1,255500535 | -0,099781133 |
| Ruminococcus E bromii B | hMGS.00005 | -6,397007584 | -1,295377642 | -0,073697343 |
| Fusicatenibacter saccharivorans | hMGS.00011 | -5,508770483 | -0,552136908 | -0,128681084 |
| Blautia A wexlerae | hMGS.00013 | -5,30561048 | -0,779105741 | -0,128321821 |
| Alistipes putredinis | hMGS.00029 | -6,795412275 | -1,010400574 | -0,054749065 |
| Dialister invisus | hMGS.00096 | -6,149496066 | -0,584502315 | -0,087525468 |
| Lactobacillus iners | hMGS.00136 | -6,23548018 | -4,344616839 | -0,057627887 |
| Blautia A obeum | hMGS.00137 | -4,642543521 | -0,216848118 | -0,078509682 |
| Prevotella bivia | hMGS.00191 | -1,962537989 | -0,418458098 | -0,407909582 |
| Fannyhessea vaginae | hMGS.00196 | -5,136095717 | -2,592986816 | -0,0965118 |
| Lactobacillus crispatus | hMGS.00239 | -9,539168818 | -5,822746644 | -0,082122963 |
| Bifidobacterium vaginale | hMGS.00248 | -4,971432285 | -2,558070883 | -0,067223862 |
| Alloscardovia omnicolens | hMGS.00407 | -4,301830082 | -0,824324435 | -0,146282752 |
| Dorea formicigenerans | hMGS.00417 | -7,035716148 | -1,123089925 | -0,069763769 |
| Streptococcus anginosus | hMGS.00468 | -2,712179593 | -0,780457497 | -0,397800869 |
| Dialister sp001553355 | hMGS.00496 | -3,758073326 | -1,043292179 | -0,42253067 |
| Dialister micraerophilus | hMGS.00576 | -3,013381796 | -1,048589413 | -0,726119294 |
| Peptostreptococcus anaerobius | hMGS.00580 | -3,392679526 | -0,446414196 | -0,40423139 |
| Aerococcus christensenii | hMGS.00589 | -4,247922223 | -1,359838233 | -0,066735952 |
| Levyella massiliensis | hMGS.00726 | -1,666359058 | -0,303967931 | -0,562424378 |
| Peptoniphilus A harei A | hMGS.00790 | -0,988095433 | -0,768880753 | -0,4962991 |
| Dialister sp900541485 | hMGS.00817 | -1,771719855 | -0,739255115 | -0,69539157 |
| Stomatobaculum sp002892395 | hMGS.00834 | -3,412242361 | -0,504587671 | -0,169391462 |
| Anaerococcus tetradius | hMGS.00902 | -3,763770053 | -1,529341099 | -0,029934306 |
| Anaerococcus vaginalis B | hMGS.00945 | -3,803198171 | -0,983975535 | -0,55542423 |
| Gleimia europaea A | hMGS.01062 | -5,772394825 | -1,366917935 | -0,585840885 |
| Peptoniphilus A lacydonensis | hMGS.01074 | -1,173529587 | -0,29672578 | -0,12534166 |
| Peptoniphilus C coxii | hMGS.01075 | -2,133447579 | -0,72022794 | -0,560548047 |
| Corynebacterium aurimucosum E | hMGS.01113 | -2,662999026 | -0,826880608 | -0,255351592 |
| Lactobacillus gasseri | hMGS.01126 | -9,734954576 | -4,336053138 | -0,044939159 |
| Limosilactobacillus portuensis | hMGS.01127 | -10,22865772 | -3,097957913 | -0,020764399 |
| Corynebacterium simulans | hMGS.01240 | -2,376772472 | -0,632982495 | -0,224380411 |
| Corynebacterium tuscaniense | hMGS.01241 | -10,5898743 | -1,13499947 | -0,211666251 |
| Ezakiella sp900540185 | hMGS.01327 | -4,809544737 | -1,153007402 | -0,26875256 |
| Finegoldia magna H | hMGS.01330 | -1,027416405 | -0,507894323 | -0,616950274 |
| Corynebacterium coyleae | hMGS.01419 | -7,584015643 | -2,88559895 | -0,431497249 |
| Anaerococcus octavius | hMGS.01500 | -2,24621748 | -0,411892709 | -0,184766393 |
| Corynebacterium aurimucosum C | hMGS.01676 | -5,617654469 | -1,641532334 | -0,083443494 |
| S5-A14a sp900553025 | hMGS.01754 | -6,415968651 | -0,834466431 | -0,130821701 |
| Lawsonella sp018376445 | hMGS.01923 | -4,281291843 | -1,104521565 | -0,744309811 |
| Atopobium minutum | hMGS.02095 | -2,932916262 | -0,347674534 | -0,185957477 |
| Corynebacterium riegelii | hMGS.02175 | -9,746069562 | -3,565396994 | -0,111946718 |
| Trueperella bernardiae | hMGS.02361 | -6,728861303 | -1,149366087 | -0,054371593 |
| Anaerococcus hydrogenalis | hMGS.02513 | -1,77823785 | -0,358225496 | -0,228327324 |
| Brevibacterium sp001815905 | hMGS.02552 | -8,039395525 | -2,120815433 | -0,107814889 |
| UBA4285 sp900542465 | hMGS.02977 | -4,561341003 | -1,319765928 | -0,687038051 |
| Varibaculum cambriense B | hMGS.03234 | -5,210681812 | -0,765936738 | -0,285433048 |
| Anaerococcus provencensis | hMGS.03321 | -2,897528717 | -0,59176277 | -0,176877111 |
| Anaerococcus vaginimassiliensis | hMGS.03324 | -3,433488526 | -0,610990504 | -0,486854582 |
| Corynebacterium aurimucosum | hMGS.03743 | -7,717204973 | -2,154055446 | -0,310991801 |
| Corynebacterium sp001767255 | hMGS.03760 | -3,353367126 | -1,447230195 | -0,817416971 |
| Corynebacterium hMGS.04874 | hMGS.04874 | -11,89020615 | -2,478582768 | -0,05917283 |
| Helcococcus hMGS.05144 | hMGS.05144 | -5,569838565 | -0,929905968 | -0,346800764 |
| JABCPO02 hMGS.05190 | hMGS.05190 | -3,701953965 | -0,699434028 | -0,248185994 |
| Faecalibacterium longum | hMGS.05756 | -5,077865421 | -0,496389931 | -0,056229189 |
| Faecalibacterium duncaniae | hMGS.05764 | -6,083463726 | -1,222320388 | -0,063492169 |
| Corynebacterium amycolatum hMGS.05820 | hMGS.05820 | -4,581610312 | -1,316319755 | -0,289153772 |
| Corynebacterium sp943913645 | hMGS.05841 | -10,74300375 | -3,517856094 | -0,522289644 |
| Anaerococcus hMGS.05963 | hMGS.05963 | -6,142895492 | -1,26599736 | -0,251518229 |
| Eremococcus coleocola | hMGS.06890 | -4,491004091 | -1,22028992 | -0,050928028 |
| Pseudoglutamicibacter albus | hMGS.07111 | -14,64027975 | -3,523842389 | -0,074477194 |
| Anaerococcus sp943912825 | hMGS.07406 | -2,087507095 | -0,891117956 | -0,334658368 |
| Fastidiosipila sanguinis | hMGS.07481 | -6,047994401 | -0,873775853 | -0,421149848 |

**Table S5:** Agreement between the three differential abundance methods (DESeq2, ANCOMBC, ALDEx2) of species consistent and significantly associated with males (higher abundance in males). FC = Fold change.

| **Species** | **MGS** | **FC_DESeq2** | **FC_ANCOMBC** | **FC_ALDEx2** |
| --- | --- | --- | --- | --- |
| Staphylococcus hominis | hMGS.00145 | 0,802425 | 0,456484197 | 0,226995081 |
| Micrococcus luteus | hMGS.00154 | 1,592029939 | 0,842401681 | 0,397151869 |
| Cutibacterium acnes | hMGS.00201 | 1,934429053 | 1,439126707 | 0,71137004 |
| Enterobacter hormaechei A | hMGS.00243 | 1,226957218 | 0,955029758 | 0,238317676 |
| Salmonella enterica | hMGS.00281 | 1,874620422 | 0,735948912 | 0,160747325 |
| Negativicoccus succinicivorans | hMGS.00519 | 2,123412184 | 1,22018649 | 0,179058302 |
| Pasteurella multocida | hMGS.00521 | 1,375014177 | 0,716173469 | 0,207926523 |
| Haemophilus seminalis | hMGS.00648 | 3,173228153 | 0,500056061 | 0,156449825 |
| Neisseria suis | hMGS.00652 | 1,959169238 | 1,233835998 | 0,513895495 |
| Enterococcus D gallinarum | hMGS.00682 | 1,952252001 | 1,283296391 | 0,586946894 |
| Aeromonas caviae | hMGS.00867 | 1,631408675 | 0,32933199 | 0,164818682 |
| Streptococcus gallolyticus | hMGS.00937 | 1,489001674 | 0,858801339 | 0,386506168 |
| Corynebacterium pyruviciproducens | hMGS.01050 | 1,457085454 | 0,784379837 | 0,099864051 |
| Klebsiella ornithinolytica | hMGS.01184 | 1,427371465 | 0,86906268 | 0,216000829 |
| Actinotignum urinale | hMGS.01225 | 5,092461837 | 3,06470248 | 0,164797625 |
| Dolosigranulum pigrum | hMGS.01244 | 3,673464375 | 1,516370385 | 0,173852485 |
| CAMCNU01 hMGS.01291 | hMGS.01291 | 3,464713926 | 2,821870878 | 0,08632074 |
| Acinetobacter johnsonii | hMGS.01295 | 2,534155612 | 0,839609028 | 0,360720239 |
| Corynebacterium glucuronolyticum | hMGS.01318 | 5,010726109 | 1,995311814 | 0,404389781 |
| Corynebacterium mucifaciens | hMGS.01537 | 1,794951576 | 0,807087424 | 0,194711036 |
| Kaistella haifensis | hMGS.01562 | 2,108154795 | 0,883176572 | 0,191707383 |
| Cutibacterium namnetense | hMGS.01684 | 1,916422229 | 0,786057222 | 0,298484596 |
| Propionibacteriaceae hMGS.01792 | hMGS.01792 | 1,951579819 | 1,439367873 | 0,623738903 |
| Brevibacterium sp005280295 | hMGS.01832 | 4,530463817 | 1,489886468 | 0,171299265 |
| Corynebacterium otitidis | hMGS.01878 | 2,050073285 | 0,664650409 | 0,126599103 |
| Enterococcus A malodoratus | hMGS.01889 | 1,871999636 | 0,980036495 | 0,228634697 |
| Mammaliicoccus sciuri | hMGS.01930 | 1,131598528 | 1,177209949 | 0,222257163 |
| Corynebacterium propinquum | hMGS.02174 | 3,627877084 | 1,598523214 | 0,150004278 |
| Latilactobacillus sakei | hMGS.02234 | 1,667610098 | 0,847948557 | 0,164022463 |
| Corynebacterium hMGS.02403 | hMGS.02403 | 5,831644499 | 1,088253926 | 0,190258686 |
| Anaerococcus jeddahensis | hMGS.02514 | 2,605717142 | 1,678786136 | 0,053413128 |
| Bacteroides sp900766195 | hMGS.02533 | 1,667481157 | 0,838726121 | 0,270109277 |
| Cutibacterium modestum | hMGS.02660 | 1,388503686 | 0,64527072 | 0,201520597 |
| Nanoperiomorbus sp004136275 | hMGS.02829 | 5,090161925 | 0,937523666 | 0,136221859 |
| Staphylococcus borealis | hMGS.02924 | 4,045574502 | 1,552404113 | 0,146681808 |
| Staphylococcus saccharolyticus | hMGS.02927 | 3,260484862 | 1,078970876 | 0,253872864 |
| Streptococcus mitis BF | hMGS.02938 | 5,791229223 | 1,426218134 | 0,174271987 |
| Citricoccus terreus | hMGS.03556 | 2,802268017 | 0,909172574 | 0,170314927 |
| Corynebacterium falsenii | hMGS.03746 | 1,755682307 | 1,280473031 | 0,353610572 |
| Lawsonella clevelandensis A | hMGS.04022 | 2,165327017 | 1,141678312 | 0,599672179 |
| Micrococcus endophyticus | hMGS.04084 | 1,800151164 | 0,926400344 | 0,199178957 |
| Staphylococcus cohnii | hMGS.04392 | 2,183935603 | 1,195198312 | 0,168762401 |
| Peptoniphilus A hMGS.05138 | hMGS.05138 | 7,289674623 | 3,12700778 | 0,185291663 |
| Mycoplasmopsis hMGS.05153 | hMGS.05153 | 3,408929184 | 2,20676102 | 0,129890392 |
| Varibaculum hMGS.05156 | hMGS.05156 | 3,76267552 | 2,1345077 | 0,203606997 |
| Propionimicrobium hMGS.05166 | hMGS.05166 | 7,791312716 | 4,338959508 | 0,158863285 |
| JAEZZM01 hMGS.05269 | hMGS.05269 | 3,816022901 | 1,512742698 | 0,133046505 |
| Anaerococcus hMGS.05274 | hMGS.05274 | 1,4831901 | 0,862201054 | 0,083124024 |
| Streptococcus suis AA | hMGS.05935 | 1,984903579 | 0,999310161 | 0,471256149 |
| Pseudomonas E oleovorans | hMGS.05946 | 2,929268435 | 0,615461147 | 0,205912684 |
| Corynebacterium gottingense | hMGS.06055 | 6,803361611 | 1,820450638 | 0,320830514 |
| Moraxella a cinereus hMGS.06075 | hMGS.06075 | 2,4543194 | 0,294613429 | 0,270918166 |
| QFNR01 sp943914555 | hMGS.06125 | 1,789735458 | 1,480742238 | 0,179274716 |
| Afipia sp000497575 | hMGS.06161 | 3,217299525 | 1,191355565 | 0,105323827 |
| Psychrobacter sanguinis | hMGS.06200 | 1,283828246 | 1,160682398 | 0,198269658 |
| Ochrobactrum tritici | hMGS.06273 | 1,315283018 | 1,02494454 | 0,210492961 |
| Halomonas stevensii | hMGS.06317 | 2,483170582 | 1,097495903 | 0,433966735 |
| Cloacibacterium normanense | hMGS.06447 | 0,695806632 | 0,72957755 | 0,260098653 |
| Luteimonas D sp002324875 | hMGS.06588 | 5,374076923 | 1,46116297 | 0,188310826 |
| Dermabacter vaginalis | hMGS.06607 | 6,910339681 | 1,445583809 | 0,268836638 |
| Corynebacterium suicordis | hMGS.06622 | 2,095573493 | 0,710958665 | 0,208985639 |
| Micrococcus sp023155805 | hMGS.06899 | 2,643715471 | 0,922756644 | 0,224199679 |
| Corynebacterium testudinoris | hMGS.07250 | 0,983583926 | 1,158698734 | 0,230965297 |
| Stutzerimonas stutzeri | hMGS.07298 | 2,515125214 | 0,98334827 | 0,246940552 |
| Corynebacterium sp943912665 | hMGS.07606 | 6,077079719 | 1,629617058 | 0,428554762 |
| Malassezia globosa | hMGS.10047 | 2,304829185 | 1,061242469 | 0,284526661 |
| Malassezia restricta | hMGS.10057 | 1,770509665 | 1,242584802 | 0,373758548 |
